# Diverse high-fat diets drive multi-omic reprogramming that persists after dietary reversal

**DOI:** 10.64898/2026.03.17.708620

**Authors:** Andrew G Van Camp, Jiwoon Park, Elif Ozcelik, Onur Eskiocak, Kadir A Ozler, Katie Papciak, Santhilal Subhash, Hanan Alwaseem, Ilgin Ergin, Charlie Chung, Vyom Shah, Brian Yueh, Aybuke Alici, Miriam R Fein, Ceyda Durmaz, Christopher Mozsary, Ece Kilic, Namita Damle, Deena Najjar, Theodore M Nelson, Krista A Ryon, Daniel J Butler, Chirag J Patel, Christoph A Thaiss, Kivanc Birsoy, Christopher E Mason, Cem Meydan, Braden T Tierney, Semir Beyaz

## Abstract

Dietary fat composition modulates host physiology and the gut microbiome, but the long-term effects of specific fat sources and the extent to which these changes resolve after dietary reversal remain incompletely defined. Here, we present a longitudinal multi-omic resource of mice maintained for one year on a purified control diet, seven high-fat diets differing in predominant fat source, or reversal regimens in which animals were switched from high-fat to control diet after 4 or 9 months. We further incorporated two cohorts with distinct pre-existing microbiome configurations to determine how baseline community structure shapes diet-induced remodeling of the gut microbiome ecosystem. By integrating longitudinal phenotyping, fecal metagenomics, fecal metabolomics, plasma metabolomics and lipidomics, and intestinal single-cell RNA sequencing, we defined the shared and dietary fat-specific responses across host and microbiome compartments. Baseline microbiome composition strongly influenced microbial responses to diet, indicating that pre-existing community structure is a major determinant of dietary ecosystem remodeling. Although many altered features shifted toward baseline after dietary reversal, only approximately half of diet-associated microbial changes recovered within the study window. A subset of taxa exhibited persistent alterations, including sustained depletion of *Lactobacillus johnsonii* and *Bifidobacterium pseudolongum* and sustained enrichment of *Alistipes finegoldii*, consistent with a “microbiome memory” of prior high-fat diet exposure. This memory effect is mirrored in the host, by sustained suppression of major histocompatibility complex class II (MHC-II) gene expression in intestinal epithelial cells after dietary reversal. These findings indicate that dietary fats leave a lasting imprint on the host-microbiome interactome that survives dietary intervention. Together, these data establish a resource for defining how dietary fat source, baseline microbiome composition, and dietary history shape host–microbiome states. The entire resource is available online as an RShiny app.

## Introduction

Dietary fat is a major determinant of host physiology and the gut microbiome^1,2^. Increased fat intake has been linked to obesity, colorectal cancer, and pro-oncogenic microbial compositional shifts in both human and murine models^3–6^. However, much of the link between diet and disease stems from observational studies not designed to untangle the complexities of host-microbiome interactions^7–13^. The physiologic response to dietary nutrient intake is driven both by those nutrients themselves and by their biochemical transformations by native microbes; variation in host response to these inputs stems from an individual’s extant physiology and genetic predispositions^14^. Specific genera that have been shown to increase with a high-fat diet include *Alistipes*, *Erysipelotrichaceae*, and *Helicobacter*^1,15,16^. In turn, high-fat diets (HFDs) increase intestinal tumorigenesis via activating lipid-sensing PPAR transcription factors^17^, inflammation^18^, bile acids^19^, or evoking microbiome dysbiosis^20,21^. The relationship between diet-induced obesity and its comorbid physiologies appears to stem from a complex interplay between host diet, genetic background, native gut microbiota, and immune activation^22–24^.

A central challenge in the field is that a “high-fat diet” is often treated as a single exposure, even though dietary fats differ substantially in chain length and saturation of fatty acids. Recent work shows that these differences influence anti-tumor immunity and likely have distinct effects on host metabolism, intestinal physiology, and microbial ecosystem structure^25^. Yet, many studies evaluate only one dietary formulation, making it difficult to distinguish general consequences of high-fat feeding from responses that are specific to particular fat sources. This problem is compounded by the fact that observational studies in humans are often unable to resolve causality in the setting of correlated dietary, microbial, and host variables. Controlled comparative studies in model organisms are therefore needed to determine which effects are shared across high-fat diets, which are fat-source-specific, and which emerge only over prolonged exposure. Despite epidemiological associations, the mechanistic interplay between host health, dietary fat, intestinal cell networks, and the microbiome remains poorly understood, in large part due to the inherent complexity of the diet-microbe-host axis^10^.

Another unresolved issue is reversibility. Early work showed that diet-induced obesity is associated with marked but partially reversible remodeling of the mouse gut microbiome^9^, while later studies demonstrated that prior dietary exposure can leave persistent microbiome alterations that shape subsequent metabolic responses even after dietary reversal and weight loss^26^. These observations raise the possibility that dietary history functions as a biological memory. However, the degree to which such persistence generalizes across diverse fat sources remains unclear. It is also not well understood whether this putative ecological memory is mirrored in host tissue programs. This question is particularly important in light of evidence that a lard-based high-fat diet suppresses major histocompatibility complex class II (MHC-II) expression in intestinal epithelial cells by triggering microbiome dysbiosis and thereby perturbs epithelial immune surveillance and promotes tumor initiation^20^.

A further source of complexity is baseline microbiome composition. Pre-existing community structure can shape how the gut ecosystem responds to perturbation, and even closely related hosts may exhibit divergent microbial and host outcomes when exposed to the same intervention^27^. Additionally, baseline microbiota can feasibly alter response to dietary input, leading to variation in host response^28,29^. This means that conclusions drawn from a single specific-pathogen-free microbial configuration may not generalize across cohorts or facilities. Comparative designs that explicitly incorporate distinct baseline microbiomes are therefore essential for identifying robust dietary signatures and for separating context-dependent effects from broadly conserved responses.

Here, we generated a longitudinal multi-omic resource to define how diverse dietary fats remodel the host–microbiome interface across time and after dietary reversal. We profiled mice maintained for one year on a purified control diet, seven high-fat diets differing in predominant fat source, or reversal regimens in which animals were returned to control diet after 4 or 9 months. To assess the contribution of baseline microbial ecology, we further incorporated two cohorts with distinct pre-existing microbiome configurations. We integrated longitudinal phenotyping with fecal metagenomics, fecal metabolomics, plasma metabolomics and lipidomics, and intestinal single-cell RNA sequencing. This design enables comparison of shared and fat-specific responses across host and microbiome compartments, identification of features that persist after dietary reversal, and evaluation of how baseline microbiome composition shapes diet-induced ecosystem remodeling. By making these data available as an interactive resource, this study provides a framework for investigating how dietary fat source and dietary history leave lasting imprints on gut microbial ecology and intestinal host state. This resource is available in an RShiny app at https://btierneyshiny.shinyapps.io/high-fat-diets-multi-omics/.

## Results

### Systematic measurement of the impact of isocaloric, high-fat diets on host physiology

To systematically assess the functional significance of different dietary lipids on the host and microbiome, we designed custom isocaloric HFDs (Envigo, 4.6 kcal/g total, 22.7% fat by weight, 45% kcal from fat) that have identical macronutrient (protein, carbohydrate and fat) and micronutrient (vitamins and minerals) composition with different fat sources and a low-fat purified control diet (Envigo, 3.8 kcal/g, 7.2% fat by weight and 17% kcal from fat, soybean oil-based) (Fig 1a, Supp Table 1). We chose fat sources that are enriched in major fatty acid types (saturated, monounsaturated and polyunsaturated fatty acids) from plants and animals as follows: lard (enriched in PA and OA), coconut oil (enriched in stearic acid (SA) and myristic acid (MA)), milkfat (enriched in PA, OA and short-chain fatty acids such as butyric acid (BA)), palm oil (enriched in PA), olive oil (enriched in OA) and fish oil (enriched in eicosapentaenoic acid (EPA) and docosahexaenoic acid (DHA)) (Fig 1b). In addition, we designed a lard-based ketogenic diet with very high fat and low carbohydrate abundance (Envigo, 7.0 kcal/g, 70.2% fat by weight, 90.3% kcal from fat and fat to protein and carbohydrate ratio is approximately 4:1). We used 8-week-old C57BL/6J (B6) mice (n=20 mice (10 male and 10 female per diet) from Jackson Laboratory and first acclimated them for one week in the animal facility, feeding all mice a purified control diet for two weeks and after that started feeding them with the purified diets described above. We performed longitudinal monitoring for up to 12 months (Fig 1c) to determine organismal metabolic alterations such as changes in body weight, glucose tolerance and metabolic cage measurements including respiratory exchange ratio (RER). This was completed in two independent cohorts that contained different initial microbiome compositions.

**Figure 1.**
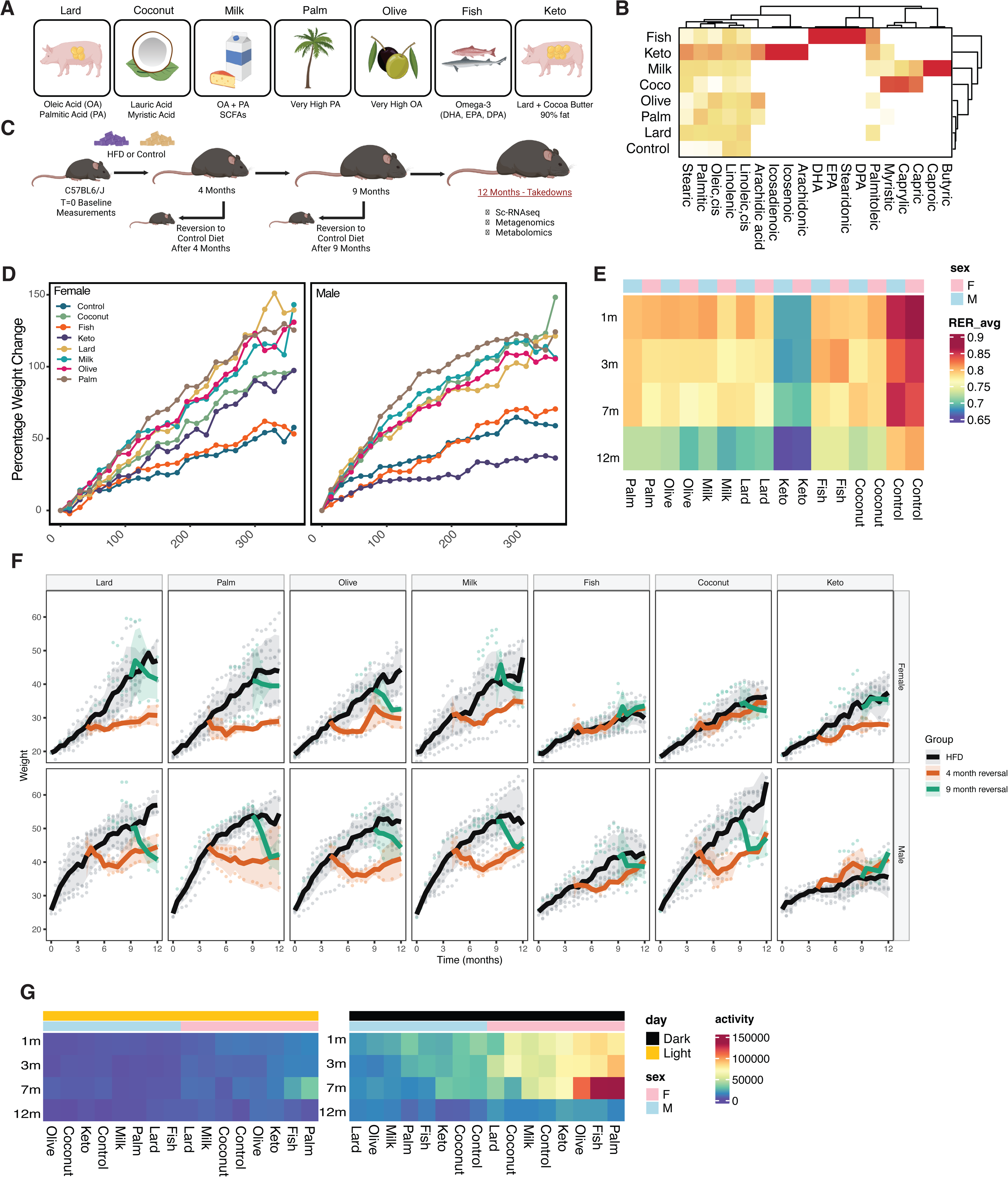
Systematic measurement of the impact of isocaloric high-fat diets on host physiology. A. Longitudinal study design in C57BL/6J mice. Subsets of animals were reverted to control diet after 4 months or 9 months on HFD, and all cohorts were terminated at 12 months for multi-omic interrogation (microbiome profiling, metabolomics, and scRNA-seq). Schematic created with BioRender. B. Heatmap of dietary fatty-acid composition across diets. Values represent the relative abundance of each fatty acid across all diets, normalized by column. Hierarchical clustering groups diets and fatty acids by similarity. C. Summary of distinguishing lipid features of each diet/fat source. Schematic created using BioRender. D. Trajectories of body weight over 12 months shown as percent change from starting weight. E. Respiratory exchange ratio (RER) measured by metabolic cage at indicated timepoints (1, 3, 7, and 12 months), displayed as a heatmap by diet and sex annotation. F. Average body weight in diet-reversion cohorts, shown by diet and sex across time, with colors indicating continuous HFD, 4-month reversal, and 9-month reversal cohorts. For each diet, male cohorts included n = 8 continuous-HFD mice, n = 3 four-month-reversal mice, and n = 3 nine-month-reversal mice; female cohorts included n = 9 continuous-HFD mice, n = 2 four-month-reversal mice, and n = 3 nine-month-reversal mice. G. Locomotor activity measured by metabolic cage across light and dark cycles (day annotation), calculated as the average of X- and Y-axis infrared beam-break (laser disruption) counts, shown longitudinally by diet and sex.

We additionally separated mice into “Reversal” (R) and “Non-Reversal” (NR) arms. Non-reversal mice received a given diet for the entire 12 month duration of the study. Reversal mice received diets for only a portion of the study before being reverted back to the control diet. The purpose in doing so was to evaluate longer-term changes to mouse physiology (e.g., permanent microbial species loss) as a function of diet. Mice were reverted from HFDs to the control diet at either four months (4MR) (3 male and 2 female mice per diet) or at the nine month sampling (9MR) (3 male and 3 female mice per diet).

Mice experience discrete phenotypic alterations as a function of sex and diet. The average weight of all diet-sex groups increased by the end of the study (Fig 1d). Relative to the control, all high-fat diets appeared to be increased except for fish oil, which roughly tracked with the control, and keto, which resulted in more weight gain in female mice and less weight gain than the controls in male mice. RER decreased with time, more so for the high-fat diets, particularly keto, compared to the control (Fig 1e). Reversions tended to lose weight after being put back on the control diet (Fig 1f), except for keto, for which males gained weight after reversion, indicating potential lingering effects of dietary intervention. In males, diet had a limited impact on activity level, while in females, palm and fish oil in particular were associated with increased activity levels (Fig 1g).

### Disentangling sex, diet, and temporal effects on microbiome diversity

First, we quantified changes in microbiome alpha diversity as a function of diet, time, and host sex in the non-reversal cohort (Fig 2A). All diets except coconut oil increased diversity relative to controls by the study’s end; however, we noted sex-specific effects in these changes. For example, by study completion, coconut oil resulted in a 1.6 fold increase in male over female mice, milkfat yielded a 1.1 fold increase, lard yielded an 1.3 fold increase, and palm oil yielded an 1.1 fold increase. The ketogenic diet induced a depletion of diversity relative to controls, more so in male mice than in females (0.9 fold decrease in male vs female mice by study completion). Olive oil induced increases in both male and female mice continuously throughout the study period. Other diets, such as palm oil and milkfat, experienced increases at the beginning that leveled off towards the end, indicating variable time-dependent impacts on overall community composition.

**Figure 2.**
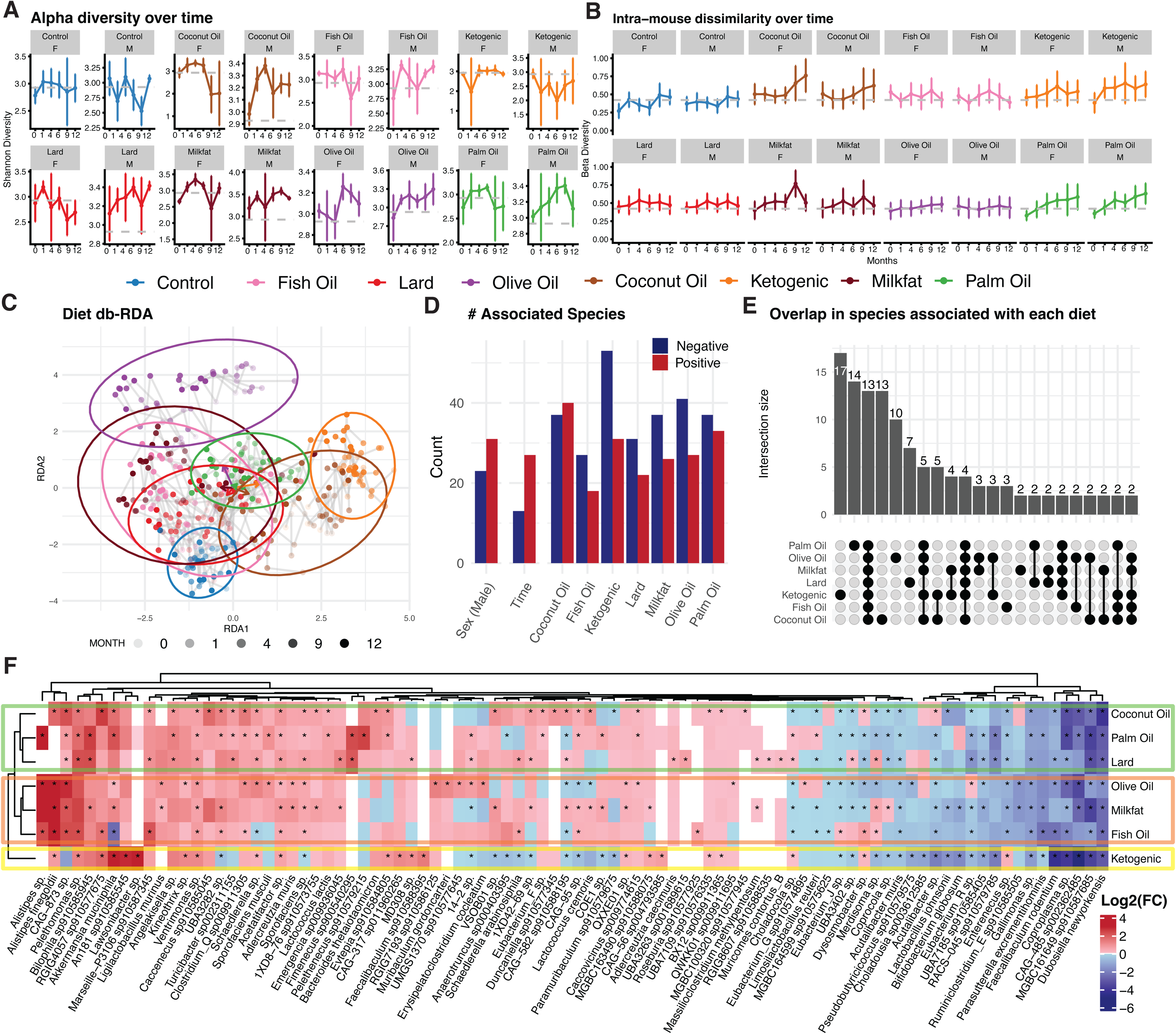
Associations between high-fat diets and gut microbiome composition. Diet colors are shared under panels A and B. A. Shannon diversity across diets and time by sex (n = 71 mice, 376 longitudinal fecal samples; non-reversal arm). X axis, month of sampling; Y axis, diversity. F = female, M = male. Vertical bars, standard deviation within each diet–time–sex group. B. Beta diversity across diets and time by sex (same mice and samples as panel A). Axes and symbols as in panel A. C. Distance-based redundancy (db-RDA) of overall microbiome composition constrained by diet (same mice and samples as panel A). Each point is one fecal sample from one mouse; lighter points are earlier in the study, darker points later. Ellipses summarize where most samples per diet fall (approximate 95% confidence regions). Grey lines link the same animal across timepoints. Arrows show how strongly each diet aligns with the ordination axes. D. Counts of species passing Benjamini–Hochberg FDR < 0.05 in generalized linear models of abundance versus metadata, after requiring the same direction of association at more than one timepoint and at most one high-fat diet association per species (sex-by-diet interaction panel omitted). Bars split by model term: sex, linear time, and diet versus control; blue, positive association; red, negative. Further filtering is described in Methods. E. UpSet plot of diet-associated species using the same significance and persistence rules as panel D. Each column is a diet; dots mark which diets share an intersection; bar height is the number of species; intersections with fewer than two species are omitted. F. Heatmap of mean log2 fold change of each species’ abundance on each high-fat diet versus control (pseudocount in the ratio), for the species set from panel D. Asterisks match the FDR rule in panel D. Species labeled only in supplementary tables are omitted from the axis. Colored outline boxes are manual diet groupings.

We next studied changes in beta diversity as a function of diet and mouse sex (Fig 2B), aiming to find if 1) a single diet induced consistent shifts in microbiome content over time or if 2) the same diet induced distinct shifts across time. Overall, we observed limited differences between sexes. Coconut oil, palm oil, and the ketogenic diet had high degrees of individual-specific-effects, with within-diet beta diversity increasing as the study continued. By comparison, though they had dramatic effects in alpha diversity, olive oil, fish oil, and lard had limited changes in beta diversity compared to the initial timepoint, indicating a consistent effect on overall microbiome composition independent of time. Together, these results indicate that sex has a distinct effect on the murine microbiome compared to high-fat dietary interventions.

### The impacts of high-fat diets on microbial community composition

These dramatic and temporally variable changes in diversity motivated a deeper analysis of species-level microbiome variation between diets. Dimensionality reduction (Fig 2C) characterized the degree to which treatment groups explained variation in gut composition versus the control diet (Fig 2C, blue circle). Plotting 95% confidence intervals for each diet indicated that, overall, the control group had the most consistency across time within subjects. Olive oil, palm oil, and the ketogenic diet demonstrated the greatest divergence from the controls. The confidence intervals for lard, fish oil, and coconut oil overlapped, demonstrating potential similarity in microbiome composition after treatment. However, the longer treatment continued (points with increasing opacity in Fig 2C), the more dissimilar from the control mice these mice appeared.

Via linear modeling, we identified species-level associations with individual diets, sex, and time (see *Methods*) as compared to the control mice. We defined statistical significance as a False-Discovery-Rate adjusted p-value (i.e., q-value) of less than 0.05, and an association between a given microbe and a diet was considered conserved if it was significant and consistent in direction across at least two timepoints. We only considered stool samples for this analysis, though the full data resource contains metagenomic samples from the entire digestive tract.

High-fat diets depleted more species than they enriched (Fig 2D). The ketogenic diet had the most substantial effects on the host gut (significantly increasing the abundance of 53 species and decreasing the abundance of 31), followed by coconut oil, palm oil, and olive oil. This effect of ketogenic diet is likely due to its difference from the isocaloric high-fat diet panel in both caloric density and macronutrient composition. Dietary associations overwhelmed sex effects (23 species with positive associations with male mice, 31 negative) and temporal (i.e., aging) effects (27 linearly increasing, 13 linearly decreasing).

Across all treatments, a total of 110 unique species had significant and consistent positive associations with diet, and 61 had analogously negative associations. We observed a greater conservation across all high-fat diets in negative-direction compositional effects as compared to enriching effects. That is, high-fat diets reduced the abundance of similar species but enriched different ones. Ketogenic, coconut oil, and palm oil had the most distinct associations, with 19, 16, and 14 unique and significant associations, respectively (Fig 2E). Thirteen species had a significant association with all diets. Twelve of these decreased in abundance relative to controls: *Acutalibacter muris, Acutalibacter sp910587835, CAG-582 sp910588195, Coproplasma sp013316055, Dubosiella newyorkensis, Enterenecus sp910585265, Eubacterium_J sp910579075, Gallimonas sp910585595, Merdisoma sp009917555, MGBC161649 sp910587685, UBA3263 sp001689615,* and *Ventrimonas sp003669055*. No species were universally enriched across diets (based on statistical significance). *Akkermansia muciniphila*, which has reported beneficial associations with numerous human diseases^30^, came close, having statistically significant, positive associations with all diets except lard (where it still underwent an albeit non-significant, but overall increase relative to controls).

Based on log2 fold changes (L2FCs) in associated species, diets hierarchically clustered into three groups (Fig 2F, left heatmap dendrogram and associated overlayed boxes): 1) coconut oil, palm oil, and lard grouped together, 2) olive oil, milkfat, and fish oil grouped together, and 3) ketogenic was separate from the rest. These groupings were characterized by numerous human-health associated species. The coconut/palm/lard group was defined by an increase in *Alistipes spp*, notably *Alistipes finegoldii. Alistipes finegoldii* is associated with modulating gut inflammation and immune responses. *Bacteroides thetaiotaomicron*, which produces short-chain fatty acids beneficial for colon health, was enriched by L2FC in all diets except milkfat and significantly enriched in coconut oil^31^. *Bifidobacterium pseudolongum subsp. globulosum,* which has anti-inflammatory effects and supports immune homeostasis^32^, was depleted in all diets except the ketogenic. *Lactococcus lactis subsp. cremoris*, which is associated with decreased cholesterol and increased glucose tolerance^33^, was increased in the coconut oil diet. *Lactococcus lactis,* distinct from *L. lactis subsp. cremoris* and associated with beneficial immunomodulatory effects^34^, was enriched significantly in the lard diet. Similarly, numerous species with potential impacts on host health were depleted by different high fats. Well-studied organisms depleted in most or all high-fat diets were *Dubosiella newyorkensis, Parasutterella excrementihominis, Faecalibaculum rodentium, Lactobacillus johnsonii,* and *Limosilactobacillus reuteri*. Many of these species, notably *F. rodentium* (and its human homologue, *Holdemanella biformis*) and *L. johnsonnii*, are involved in maintaining gut homeostasis and exhibit anti-tumor effects^35–40^. Finally, we noted that the gut pathobiont *Clostridium difficile* was notably enriched in olive oil by the end of the study.

### The longitudinal trajectories of microbial communities as a function of diet

Our initial analysis identified associations with diet, but it did not account for how species abundance might vary across time after a dietary change. As we observed with *Akkermansia muciniphila* (Fig 3A, top left) and fish oil, the relationship between time, diet, and certain microbial species was not uniformly linear. Indeed, many of the additional health-associated organisms, (e.g., *L. johnsonii*) identified in Figure 2 demonstrated variable trajectories over time (Fig 3A), with some being permanently depleted, and others (e.g., *D. newyorkensis*) appearing to recover slightly by the end of the study in all diets.

**Figure 3.**
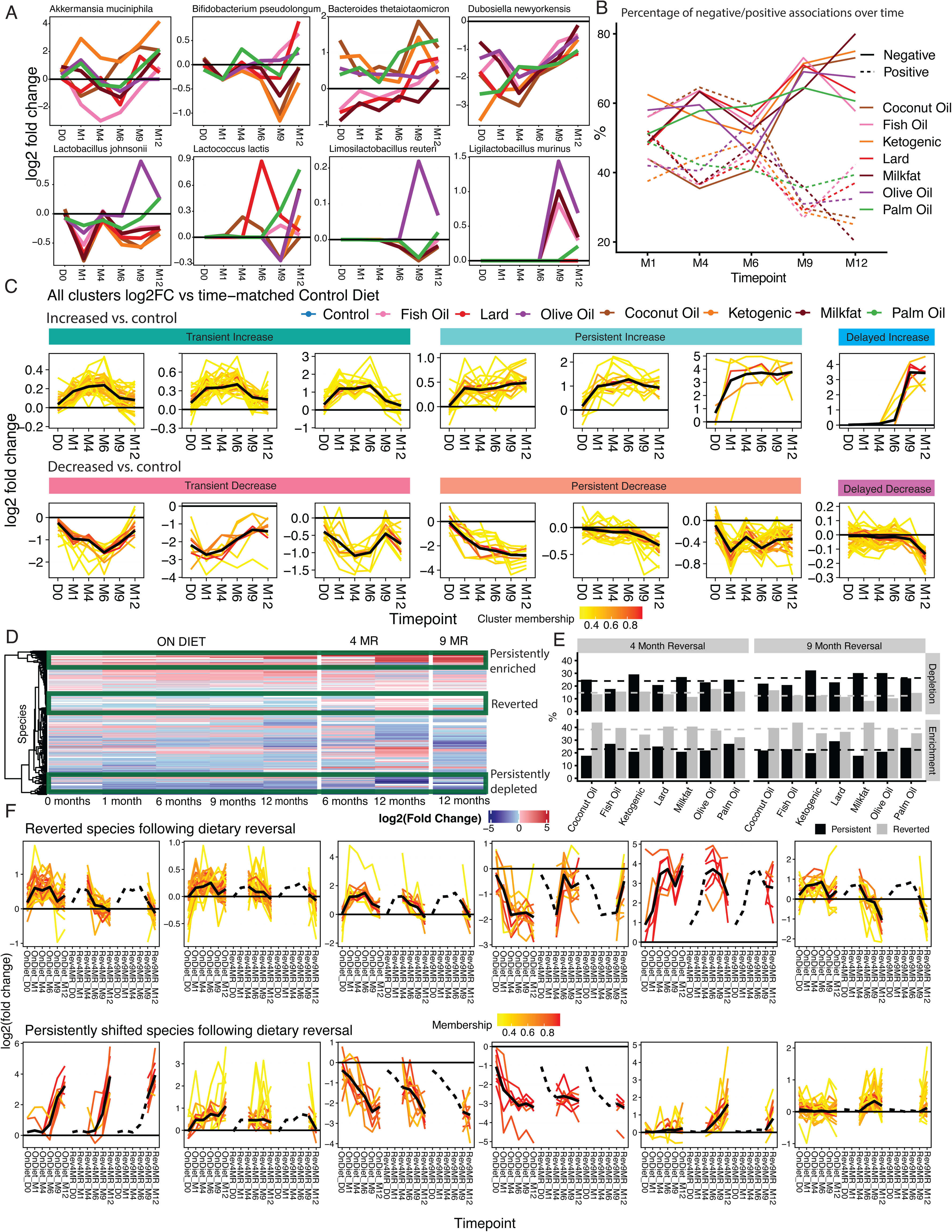
Longitudinal microbiome shifts with and without dietary reversal in the H- vs. H+ cohorts; n = 158 mice, 809 fecal samples total: non-reversal 71 mice / 376 samples; four-month reversal 38 mice / 136 samples; nine-month reversal 49 mice / 297 samples). Reversal status is defined per mouse and applies at all timepoints for that animal. Diet colors follow panel B and are reused in panels A, C, and F. A. Species-level trajectories: average log2 fold change in relative abundance versus time-matched control for each high-fat diet (n = 158 mice, 809 fecal samples). Zero line, no difference from control; horizontal axis, study month (e.g., M1, M4). B. For each diet, the fraction of associations at each timepoint (solid vs dashed; should sum to 100% at each time) (n = 158 mice, 809 fecal samples). C. Fuzzy c-means on longitudinal log2 fold change profiles for non-reversal mice (k = 20; 14 clusters shown, 6 omitted as redundant or low signal; four omitted clusters were near-flat by a post hoc center rule, max |mean log2 fold change| across on-diet times < 0.2973; two showed small non-flat variation and were dropped for clarity) (n = 71 mice, 376 fecal samples). Each thin line is one species in one mouse; color is cluster. Axes match panel A. D. Heatmap of log2 fold change versus control across reversal and non-reversal; rows are species (hierarchical clustering), columns calendar time and diet phase. Green outlines, annotated persistent enrich, revert, or deplete groups (n = 158 mice, 809 fecal samples). E. Counts of species persistently enriched or depleted after four- or nine-month reversal (one count per species per diet per reversal arm). “Persistently increased/decreased” and “reverted” as in the main text (n = 158 mice, 809 fecal samples). F. Clustered microbial trajectory profiles as in panel C but with reversal on the X axis: on-diet block (OnDiet_D0–M12), four-month reversal block (Rev4MR_…), nine-month reversal block (Rev9MR_…); dashed segments, pre-reversal timepoints in reversal arms (n = 158 mice, 809 fecal samples).

We observed that in the short term (six months into the study), high-fat diets enriched and depleted similar proportions of microorganisms (Fig 3B). However, by the study completion, the majority of lingering associations were depletionary. In other words, in the long term, organisms initially enriched by a dietary intervention decreased in abundance, whereas, organisms that were depleted tended to stay depleted, the number of lost species increasing over time, indicating that high-fat diets, overall, have a mostly repressive effect on the gut microbiome.

We observed (Figure 3C) discrete time signatures of microbial species that varied across diets. We used c-means clustering to group the longitudinal log2 fold changes (L2FC) of the organisms identified in Figure 2D-F. We were able to group within-diet species variation across time into six categories: transient increases, persistent increases, delayed increases, transient decreases, persistent decreases, and delayed decreases.

We hypothesize that the existence of these time trends could be used to disentangle complex network dynamics in response to microbial diets. For example, the initial enrichment and later depletion of a species could be a natural course correction by the gut responding, over time, to an introduced shift. Similarly, a delayed onset in abundance changes could be a latent ecological restructuring due to the continued exposure of a dietary stressor.

### High-fat diets induce changes to gut microbiome composition that persist even following cessation of dietary treatment

We next interrogated the impact of dietary reversals on gut microbiome composition. This was enabled by the two cohorts of mice that were reverted from high-fat diets back to the control diet at 4 months and 9 months into the study, respectively. We hypothesized that by comparing the longitudinal trajectories of different taxa, we could discriminate between irreversible, diet-induced changes to the gut microbiome versus those that would return to baseline with the original diet.

We observed that across all species associated with dietary shifts (Fig 3D), there were numerous cases where the relative abundance of a given organism was persistently depleted or enriched both before and after reversal, indicating long-term effects on gut microbiome composition due to dietary intervention. For both the four and nine month reversals, about half of species (53.1% and 51.3%, respectively) reverted to near baseline levels following reversion to the control diet. For the 4 month reversals, 24.0% of species observed were persistently depleted, and 22.9% were persistently enriched. For the 9 month reversals, these values were 26.3% and 22.3%, respectively (Fig 3E). Species that were enriched by high-fat diet were more likely to revert to their baseline levels following reversal (Fig 3E, bottom row, grey bars). Ketogenic, milkfat, and palm oil caused the greatest number of persistent depletions, and lard, fish oil, and palm oil resulted in the most persistently enriched species.

As with the non-reversal cohort, we used c-means clustering to interrogate the longitudinal trajectories of species before and after reversal (Fig 3F). There were discrete patterns of enrichment, depletion, and return to baseline in the raw log2FC data. These further highlighted the strength of persistent depletions, with organisms dropping by five-fold log2FC in many cases, and showing no signs of returning even following reversal. Even when a species began to return to levels similar to the control mice, it rarely returned all the way to baseline abundance during the observation window of the experiment.

Finally, we highlighted the trajectories of some of the human-health-associated species that were identified as diet-associated in Fig 2 (Supp Fig 1). These included *A. finegoldii*, *A. muciniphila, L. johnsonii, B. pseudolongum,* and *B. thetaiotaomicron*. The strong, persistent enrichment of *A. finegoldii* in multiple diets was apparent, whereas the depleting effect of the ketogenic diet (especially for *L. johnsonii* and *B. pseudolongum*) was equally clear. The ketogenic diet was also associated with persistent enrichment of *A. muciniphila* and *B. thetaiotaomicron*, although the biological consequences of these shifts cannot be inferred from the present dataset alone.

### Baseline microbiome composition drives response to dietary intervention

Gut microbiome composition acts as a filter through which dietary inputs are processed, yet most dietary studies rely on a single, standardized specific-pathogen-free (SPF) flora that may lack ecologically relevant pathobionts. The degree to which baseline microbiome composition drives response to dietary intervention is not fully known. However, it could feasibly modulate downstream ramifications on the host. For example, microbiome composition is known to modulate response to antibiotics, cancer immunotherapy, and other therapeutics^27,41–43^. Our previous work has established that a lard-based high-fat diet dampens intestinal epithelial immune surveillance and MHC-II expression in part by depleting pathobionts including *Helicobacter* species *H. typhlonius*^20^.

To rigorously test the impact of baseline microbiome composition on host and microbiome alterations, we incorporated a distinct cohort of mice harboring a natural, diverse microbiota containing the pathobiont *Helicobacter typhlonius* (H+, see Methods). 151 species were found between both cohorts, with 83 unique to the original cohort (OC), and 113 unique to the H+ cohort (Supp Fig 2). The alpha diversity in the H+ cohort tended to be higher (p < 0.05, Supp Fig 2A) than in the OC cohort across all diets except the ketogenic diet, palm oil, and milkfat. The H+ mice were notable because their gut microbiota were enriched for the family *Helicobacteraceae* (notably *Helicobacter typhlonius*) (Supp Fig 2B), which is known to be associated with tumorigenesis and MHC II expression^20,44^. Compared to the OC, the H+ cohort was additionally enriched in *Bacteroides uniformis* (associated with anti-obesity effects in mice)^45,46^, *Odoribacter* (species of which are associated with high-fat diets and glucose control)^47,48^, *Bacteroides acidifaciens* (also associated with anti-obesity effects, insulin sensitivity, and liver health)^49,50^. It was depleted in many of the health-relevant, diet-associated pieces in the OC, including *Akkermansia muciniphila* and *Dubosiella newyorkensis*.

We aimed to identify organisms that had 1) the same response to dietary intervention in both cohorts (i.e., robust to baseline microbiome composition), those that 2) had discordant responses, and 3) those that were uniquely found in the H+ cohort and diet-associated (Supp Fig 2C). The results stemming from the full association study, for both taxonomy and functions, can be found in Supp Fig 2. The majority of associations were distinct between cohorts. Across all conditions, 200 (11.7%) species were conserved, 829 (48.9%) were in distinct directions, and 668 (39.3%) were organisms uniquely found in the H+ cohort. Fish oil and lard had the least conservation between cohorts overall, whereas the ketogenic and palm oil had the most.

We identified several specific species that had similar or discrete responses to dietary treatment in both the H+ and OC (Supp Fig 2D). We define the former group as “robust to the baseline microbiome composition.” Many bacteria implicated in human health fell into this category, including *L. lactis, L. cremoris, Enterococcus faecalis* (depleted in palm oil/olive oil/coconut oil/ketogenic and increased in lard and milkfat), and *B. thetaiotaomicron*. *Clostridium spp.* additionally demonstrated conserved responses, being depleted in palm oil, olive oil, the ketogenic diet, and coconut oil, and enriched in fish oil and lard. However, some of the strongest associations from the OC were not universally replicated across all diets (Supp Fig 2E). This was notably the case for coconut oil, the ketogenic diet, and palm oil. *B. globulosum* was slightly enriched in palm oil in the OC cohort, but depleted in all the H+ cohort (and all other diets). *L. reuteri* and *L. johnsonii* also exhibited different responses to palm oil, being depleted by treatment in the OC and increased in the H+ cohort.

Finally, we interrogated microbiome changes in organisms and pathways that were specific to the H+ cohort. *Helicobacter C typhlonius* was significantly depleted by the ketogenic diet. It additionally was decreased in lard and in olive oil. It was increased by all other diets (though these associations were not FDR-significant). *Helicobacter C sp910577135*, though, was significantly depleted in all diets. Additionally, the genus *Bacteroides* was notably associated with dietary shifts. This included health-associated organisms like *Bacteroides acidifaciens* (associated with decreased BMI)^49^, which was increased significantly by the ketogenic diet and milkfat, and *Bacteroides uniformis* (a putatively beneficial short-chain fatty acid producer), which was significantly depleted by coconut oil.

### The landscape of high-fat diet-induced metabolomic and lipidomic changes

We additionally generated fecal and plasma metabolomics (polar and lipids for the latter). The processed resultant data tables as companion datasets for integration with host gene expression and microbiome composition data. We observed the strongest associations between diet and fecal metabolomics, followed by lipidomics, followed by serum metabolomics (Figure 4A). Fish oil, in particular, had a differential signature from the other diets. Ketogenic also stood apart from the others, but predominantly in the lipidomics.

**Figure 4.**
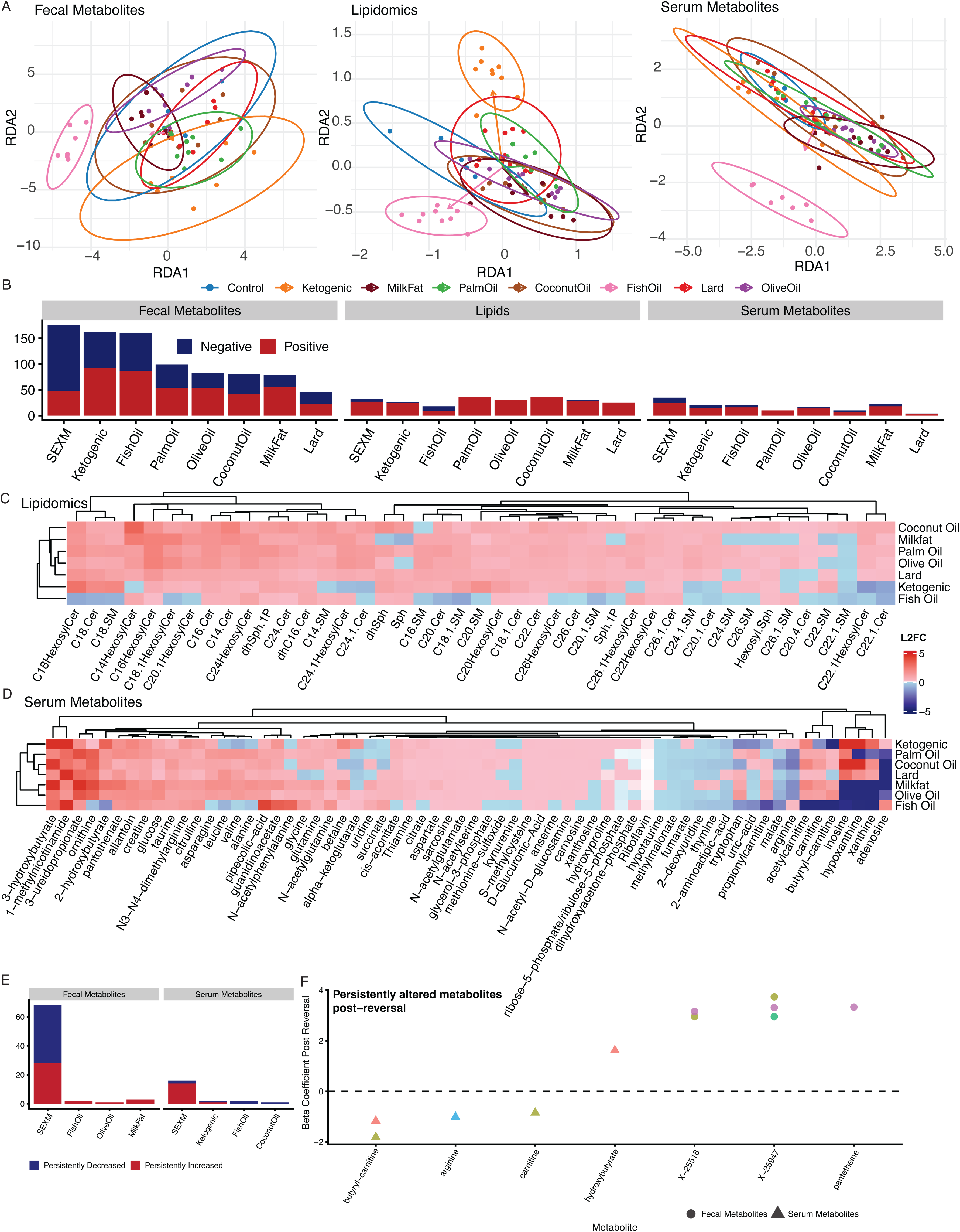
Fecal metabolites, fecal lipids, and serum metabolites by diet. Metabolite layers matched to fecal microbiome sampling at M6 and M9 (n = 71 mice with paired data). A. db-RDA per matrix (samples × log-transformed features; constraint, diet). Subpanels: untargeted fecal metabolomics M7 (n = 52 samples), fecal lipidomics M8 (n = 68), serum metabolomics M6/M11 averaged (n = 65 sample-by-sex points), each after layer-specific prevalence filtering and correlation de-redundancy where applied (Methods). B. Bar summaries of features with Benjamini–Hochberg FDR < 0.1 in linear models of log10 abundance (or normalized log measure) on sex and diet (diet and sex terms counted separately). Blue, negative association; red, positive. After filtering, models used 1048 fecal metabolites, 43 lipid species, and 94 serum metabolites. C. Lipids with FDR < 0.1 for at least one high-fat diet (n = 68 fecal lipid samples). Cell color, mean log2 fold change versus control; rows and columns clustered hierarchically. D. Serum metabolites with FDR < 0.1 (same display rules as panel C) (n = 65 serum sample-by-sex combinations as in panel A). E. Among features significant on high-fat diet and again after return to control, counts staying higher or lower (FDR < 0.1 in both windows), split by fecal metabolites, lipids, and serum. F. Features classified as persistently changed in panel E. Each point is one feature; X axis, feature name; Y axis, post-reversal association strength (Methods). Color, diet; shape, layer (fecal metabolite, lipid, serum). Horizontal dashed line, zero effect.

Again using linear modeling, we evaluated associations between these multi-omic features and host diet (Fig 4B). As with bacterial associations, the majority of fecal metabolite associations were negative. The ketogenic and fish oil diets demonstrated the greatest number of associations for diets (ketogenic: 92 positive and 70 negative; fish oil: 87 positive and 74 negative). Lard and milkfat had the fewest (lard: 23 positive, 23 negative; milkfat: 55 positive, 24 negative).

We noted that while they had fewer overall associations than fecal metabolomics, the plasma metabolite associations were almost all positive, except in the case of fish oil (Fig 4B). Plasma lipids showed increased associations with ceramides and sphingomyelins with the ketogenic diet (a hallmark of ketosis)^51^, whereas fish oil demonstrated suppression of these species compared to other fats (Fig 4C). Coconut oil demonstrated increased medium-chain hydroxylated sphingolipids, and lard and palm oil had similar signatures, especially in long-chain ceramides.

The polar metabolite data additionally yielded associations relevant to host health. The ketogenic diet increased carnitines, ketone-related metabolites (e.g., β-hydroxybutyrate analogs and acetoacetate derivatives), citric acid cycle flux, and an upregulation of amino acid catabolites. Palm oil and lard increased branched chain amino acid catabolites (e.g., xanthine), carnitines, and indicators of oxidative stress (e.g., taurine). Fish oil increased omega-3 fatty acids and, simultaneously, appeared to reduce carnitines. Coconut oil, overall, had a signature intermediate between the ketogenic diet and saturated fats, whereas milkfat had, on average, reduced association sizes but clustered with lard and palm oil.

Finally, we noted that following dietary reversal, some diet-associated metabolites were still persistently increased or decreased, mirroring the effect observed with certain taxa (Fig 4E-F). The majority of persistently altered metabolites were associated with sex; only 11 were diet associated (across fish oil, olive oil, milkfat, ketogenic, and coconut oil). Two unidentified metabolites, hydroxybutyrate, and pantetheine were increased, and three carnitines were decreased. No lipids were persistently altered according to this analysis. That being said, these values stem from rigorous False Discovery Rate adjusted cutoffs adjusted for all diet-metabolite associations; nominally significant associations yielded many more, still high effect size, persistent alterations (Supp Fig 3).

### High-fat diets alter diverse classes of fecal metabolites

We next grouped fecal metabolites into functional categories and compared their signatures across diets (Fig 5B, Supp Fig 4). As we observed in the plasma metabolomics, high-fat diets mostly affected lipid metabolism, with the ketogenic having the most outsized effects. Host sex appeared to impact lipid metabolism as well as bile acid composition and heme metabolism.

**Figure 5.**
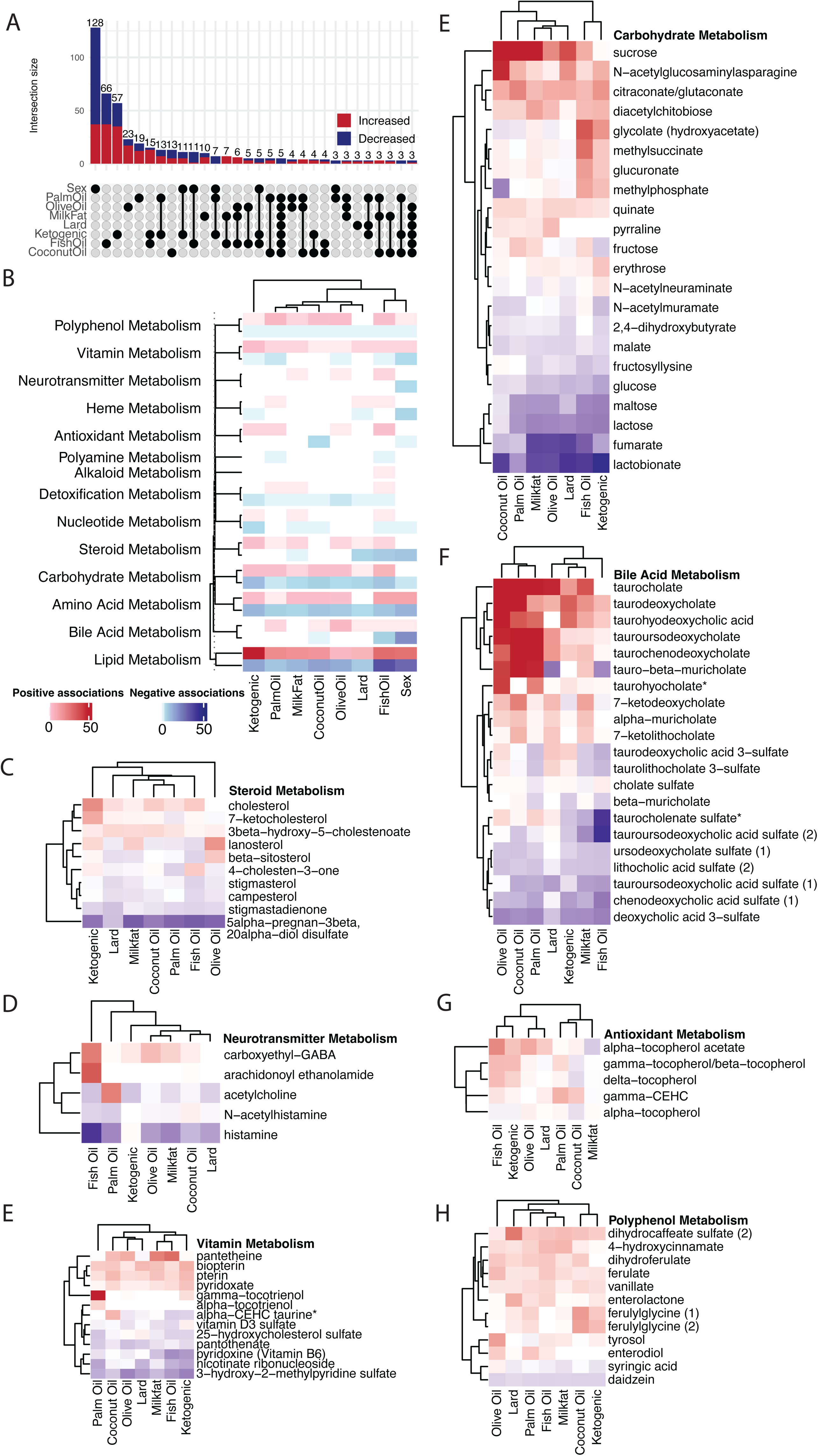
Fecal metabolite classes, diet associations, and overlaps (n = 52 samples from n = 71 mice with paired microbiome sampling; 1048 metabolite features after prevalence filtering). A. Overlap of metabolites positively or negatively associated with diet or sex at FDR < 0.05 (per-metabolite GLM, ValueLog10 ∼ sex + diet, Benjamini–Hochberg). Bar colors, direction; numbers above bars, counts. Overlap bars are descriptive (no separate test on the counts). B. FDR-significant metabolites per biochemical class by diet (n = 52 samples, 1048 features). Blue, negative associations; red, positive. C-H. Strongly diet-associated metabolites (FDR < 0.05) by class; metabolites with at least one diet association enter the heatmaps.

Fecal lipid alterations predominantly involved variation in long-chain fatty acids (Supp Fig 4A). Only a handful of short-chain fatty acids were enriched, and these were found in the ketogenic diet, coconut oil, and fish oil. Across all diets, the majority of altered lipids were saturated fats. Polyunsaturated fats were overall depleted. We observed strong variation in sphingolipids (Supp Fig 4B) and other lipid species (Supp Fig 4C). The ketogenic diet showed the most distinct profile, with marked increases in ceramides and sphingomyelins together with pronounced accumulation of acyl-carnitines and long-chain dicarboxylates, consistent with elevated β-oxidation, peroxisomal ω-oxidation, and sphingolipid turnover. Across diets, long-chain lipid responses varied substantially: ketogenic, lard, and fish-oil diets enriched very-long-chain omega-3 and omega-6 fatty acids, whereas palm, olive, and coconut oils—and to a lesser extent milkfat and lard—primarily increased palmitoyl-, stearoyl-, oleoyl-, and linoleoyl-linked species, reflecting their saturated and monounsaturated compositions. Fish oil uniquely increased EPA- and DPA-derived metabolites (e.g., 20:5n3, 21:5n3, 18:4n3), while saturated-fat diets showed stronger ceramide accumulation. Medium-chain species and dicarboxylates were broadly enriched across diets, whereas short-chain fatty acids were only modestly increased, indicating shared enhancement of fatty-acid oxidation pathways with diet-specific routing through oxidative and sphingolipid metabolism.

Additional impacted metabolite categories of interest in the context of host health included (1) steroid metabolism enriched by all diets except coconut oil and (2) neurotransmitters that were enriched by milkfat, olive oil, and fish oil. The specific steroids that were altered tended to be cholesterol and its related compounds (Fig 5C). Cholesterol had high L2FCs compared to control in the ketogenic diet, fish oil, and coconut oil. It was notably not enriched in olive oil, whereas lanosterol and beta-sitosterol were. 7-ketocholesterol, which is associated with atherosclerosis, was enriched in the ketogenic diet and depleted in palm oil and olive oil. 5alpha−pregnan−3beta, 20alpha−diol disulfate was depleted in all diets. For neurotransmitters (Fig 5D), we observed that carboxyethyl-GABA was enriched in fish oil, the ketogenic diet, coconut oil, olive oil, and milkfat. Fish oil was additionally enriched in arachidonoyl ethanolamide; all diets except the ketogenic depleted histamine. Palm oil enriched acetylcholine.

We also report diet-specific variation in carbohydrate, bile acid, antioxidant, and polyphenol metabolisms (Fig 5G-K). High-fat diets tended to universally enrich sucrose, N−acetylglucosaminylasparagine, citraconate/glutaconate, quinate, and diacetylchitobiose. Fumarate, lactose, maltose, glucose and lactobionate were depleted. Fish oil and the ketogenic diet, again similar in signature, uniquely enriched gycolate, glucuronate, methylphosphate, and methylsuccinate. The most universally enriched bile acids across all diets (though most notably in the ketogenic) were taurine-conjugated. Deoxycholic acid sulfate, chenodeoxycholic acid sulfate (1), tauroursodeoxycholic acid sulfate (1), lithocholic acid sulfate (2), and ursodeoxycholate sulfate (1) were universally depleted. Fish oil enriched the most antioxidants (gamma−CEHC, delta−tocopherol, gamma−tocopherol/beta−tocopherol, and alpha−tocopherol acetate), whereas milkfat enriched the fewest. For polyphenols, vanillate, ferulate, dihydroferulate, ferulylglycines, 4−hydroxycinnamate, and dihydrocaffeate sulfate (2) were enriched in nearly all diets, whereas daidzein was depleted. For vitamins, pyridoxate, pterin, biopterin, and pantetheine were universally enriched, and 3−hydroxy−2−methylpyridine sulfate, nicotinate ribonucleoside, pyridoxine (Vitamin B6), pantothenate, and 25−hydroxycholesterol sulfate were depleted. Gamma−tocotrienol was uniquely and highly enriched in palm oil, and vitamin D3 sulfate was uniquely enriched in the ketogenic diet.

### Fecal metabolomic alterations are correlated with diet-induced microbiome shifts

We observed alterations in some microbially-produced or associated metabolites. For example, lactobacillic acid was depleted in all diets; lactobionate and lactose were as well. This seemed reasonable, as based on the results from Figure 2, lactobacilli were reduced across most dietary conditions. To further understand the relationship between fecal metabolomics and gut microbiome composition, we used regularized regression to identify putative relationships between species composition and metabolite abundance (see *Data Availability,* FigShare repository). As an alternate, more conservative, approach, we provide the same analysis that used univariate linear regression to compute associations between metabolite-microbe pairs, enabling statistical inference and multiple hypothesis testing on each potential relationship (see *Data Availability,* FigShare repository). We found generally consistent results between both methods, though the latter, as expected, did yield fewer associations, and in the main figures we report the results from the former.

Microbial species were predominantly associated with increases and decreases lipid metabolism (Fig 6A). Bile acids were predominantly inversely correlated to different microbial species. *Alistipes finegoldii, Cholodaousia sp003612585,* and *Lawsonibacter sp910588635* had the most associated metabolites overall. Many of the species altered by diet and highlighted in Figure 2 were correlated to diverse metabolite concentrations outside of lipids alone. *A. finegoldii,* for example, was correlated to increased polyphenol metabolism, neurotransmitter metabolism, and bile acid metabolism. *L. johnsonii* was also positively correlated to polyphenol concentrations. *L. lactis* was inversely correlated with steroid levels. *A. muciniphila* was negatively correlated with neurotransmitters.

**Figure 6.**
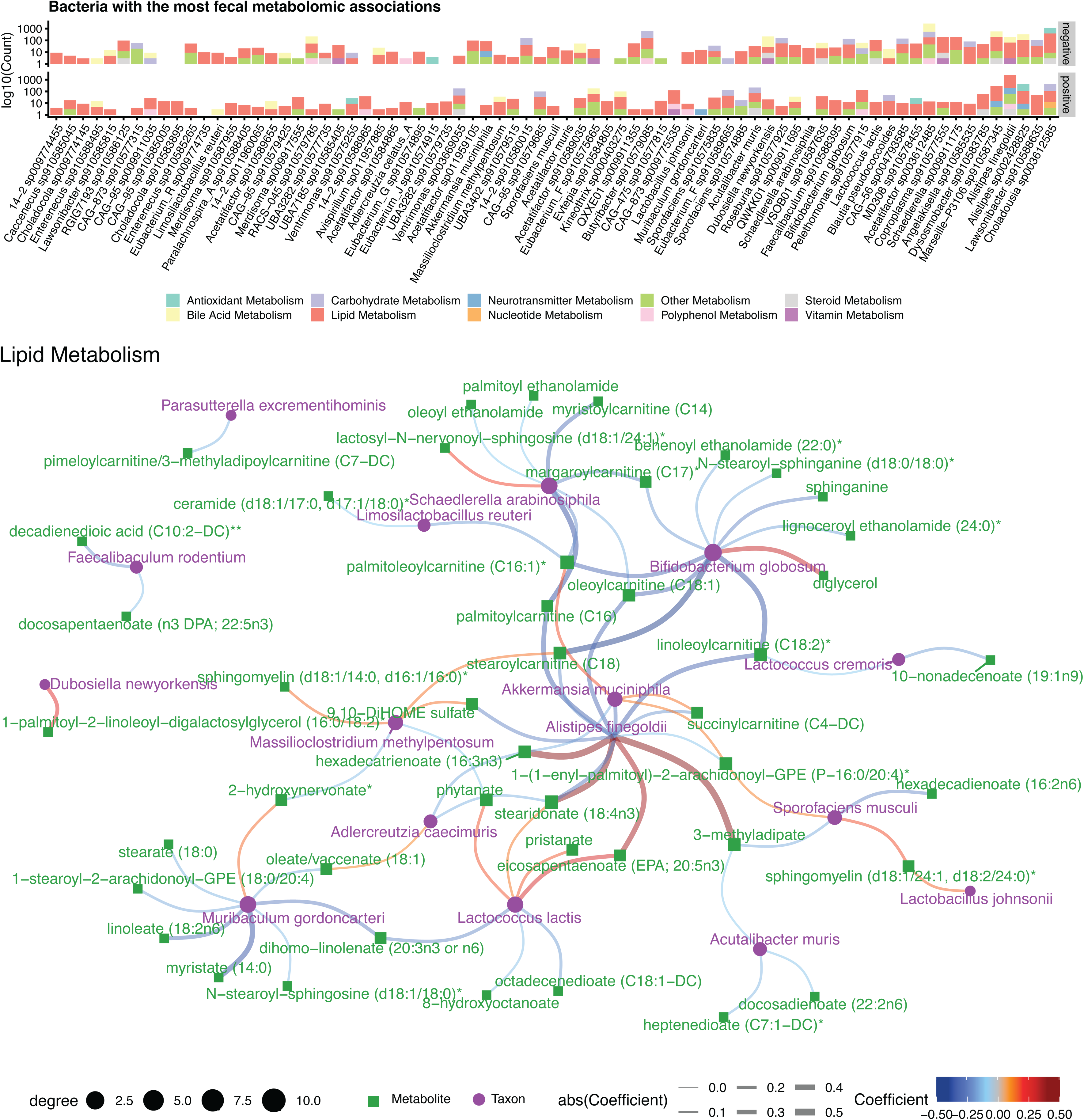
Microbe–metabolite associations (n = 71 mice and 52 fecal metabolite profiles at M7 with paired microbiome data; filtering as in Methods). A. Counts of microbial species with the largest numbers of non-zero LASSO coefficients linking their abundances to fecal metabolite levels (positive vs negative facets; point color, metabolite class). B. Network of LASSO coefficients with |coef| > 0.1 between microbes and fecal lipid-related features. Red, positive; blue, negative; line width, |coef|; green squares, metabolites; purple circles, taxa; point size, degree.

With an emphasis on well-characterized (i.e., non-candidate taxon) bacteria, we executed a network analysis to identify microbes that were potentially central to gut ecology as well as specific microbe-metabolite relationships with relevance to the host (Fig 6B). For lipid metabolism, we observed that *A. finegoldii, Schaedlerella arabinosiphila*, *A. muciniphila, B. globulosum, Limosilactobacillus reuteri,* and *Muribaculum gordoncarteri* formed a large network that included many different derivatives of stearic acid, pristanic acid, oleic acid, and palmitic acid, all of which were found in the dietary treatments in this study. In general, these organisms were inversely correlated with these fatty acids. *M. gordoncarteri* was inversely correlated with linoleate and myristate, for example. One exception to this were *Lactococcus lactis* and *A. finegoldii*, which was positively correlated to levels of pristanate, stearidonate, and phytanate. It has been reported as capable of metabolizing similar medium chain fatty acids^52^. Of additional interest are *B. globulosum* and *S. arabinosiphila*, which was negatively correlated to various ethanolamides and arylcarnatines.

Exploring networks outside of general lipid metabolism reproduced known causal relationships between microbial metabolism and the fecal metabolome. For example, *A. finegoldii* was strongly, positively correlated to carboxethyl-GABA production (Supp Fig 5A), indicating a possible direct relationship between high-fat diets and the gut-brain axis. Similarly, there were positive correlations between both *L. lactis* and arachidnyl ethanolamide, an endocannabinoid.

We additionally reproduced and identified putative novel gut microbiome relationships with steroid, vitamin, carbohydrate, bile acid, and polyphenol metabolism (Supp Fig 5B-F). For example, in accordance with the literature, *L. lactis* had a strong negative correlation with lanosterol, and *B. globulosum* had a negative correlation with 7-ketocholesterol (to which *A. finegoldii* was positively correlated). *L. cremoris* and *D. newyorkensis* were inversely associated with Vitamin B6 production. Finally, *L. reuteri* was highly associated with chenodeoxycholic acid sulfate and tauroursodeoxycholic acid sulfate.

### High-fat diets modulate host transcription of metabolic and immune genes

We analyzed differentially-expressed genes (DEGs) for each cell type observed from scRNA-seq in the mice fed different HFDs compared to the mice fed the control diet using two independent methods, DESeq2 and Wilcoxon rank sum tests. The dataset contains 20 different cell types, with 14 intestinal cell types and six immune cell types (Fig 7a). The total number of DEGs observed for each diet in each cell type via each method were plotted for cell types with greater than 100 DEGs (DESeq2 DEGs Fig 7b, Wilcoxon DEGs Supp Fig 6a). The greatest number of DESeq2 DEGs were observed in intestinal epithelial cells, including enterocytes (mature proximal and late progenitor) and goblet cells, which all had more DEGs increased compared to the control diet than decreased. In enteroendocrine cells, more decreased than increased DEGs were observed. To examine specific DEGs, the L2FC for the top five DEGs for each of the top six cell types at 12 months were plotted (DESeq2 DEGs Fig 7c, Wilcoxon DEGs Supp Fig 6b). These top DESeq2 DEGs included MHC-II genes such as H2-Aa and Cd74, as well as antimicrobial lectin genes Reg3b and Reg3g, and metabolism-related genes Slc2a2 and Cyp4a10.

**Figure 7.**
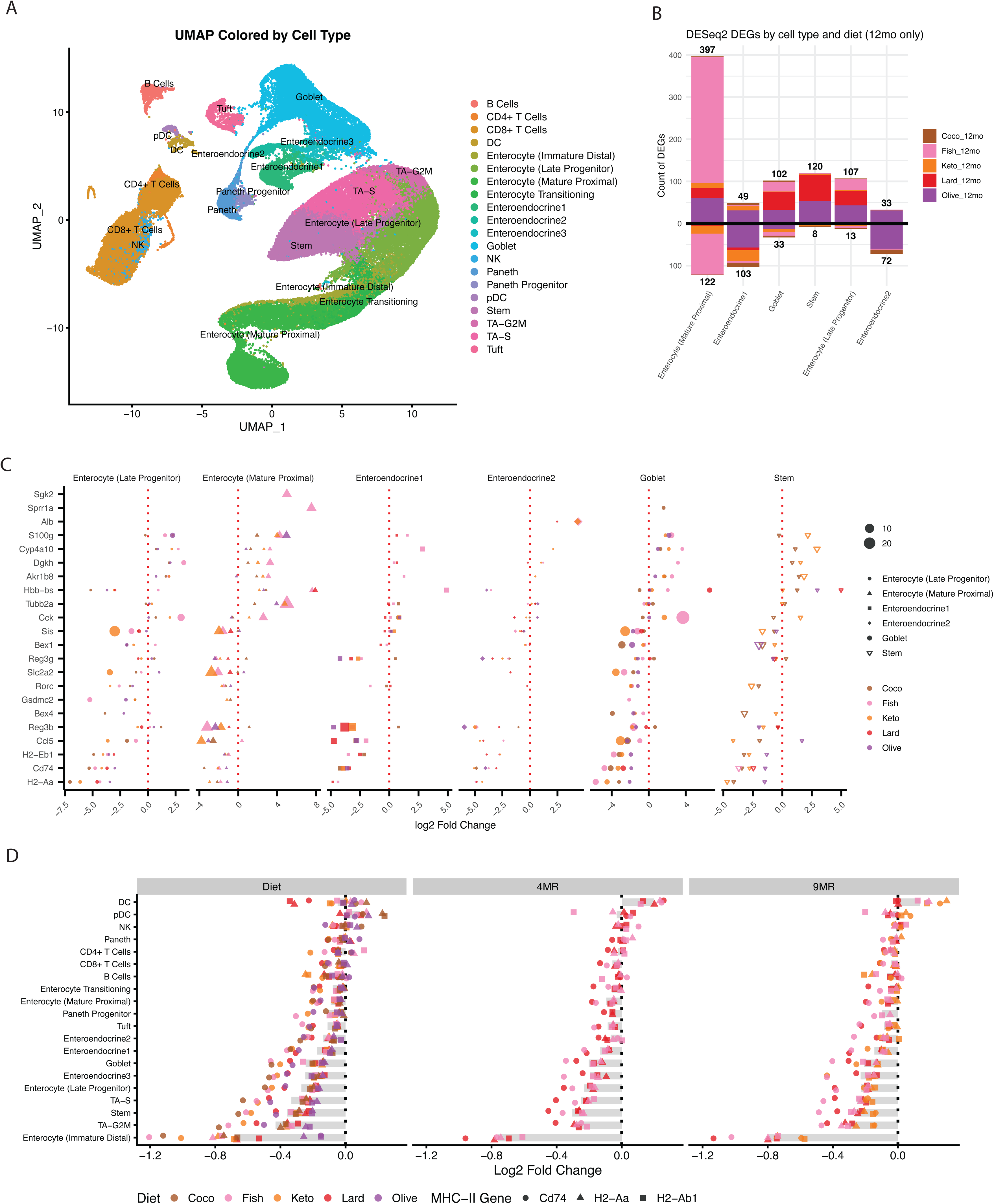
Single-cell transcriptomic profiling identifies diet-associated, cell-type-specific transcriptional changes in host tissue. A. UMAP of the scRNA-seq atlas colored by annotated cell type (n = 90,971 cells from 31 mice; 20 annotated cell types). B. Number of significantly increased and decreased differentially expressed genes (DEGs) identified by DESeq2 in male mice, grouped by cell type and diet for the 12-month diet comparisons. Bars show genes with FDR-adjusted p < 0.05. For DESeq2, raw RNA counts were pseudobulked by summing counts within each cell type for each mouse and diet, and differential expression was tested separately for each cell type and diet versus control using DESeq2 with Benjamini-Hochberg multiple-testing correction. Male 12-month comparisons included n = 3 control mice and n = 2 mice per diet group (32,895 cells total across control, Coco, Fish, Keto, Lard, and Olive groups); genes were tested after filtering for counts >= 10 in at least 2 pseudobulk samples. No additional statistical tests were applied to this summary visualization. C. Top five DESeq2 DEGs from the six cell types with the largest numbers of significant DEGs, ranked by absolute log2 fold change among genes with adjusted p < 0.05. Point position indicates log2 fold change, point size indicates −log10(FDR-adjusted p value), and color indicates diet. The underlying statistical framework, biological replicate numbers, and multiple-comparison correction are the same as in (B) (n = 3 control mice and n = 2 mice per diet group for each comparison). No additional statistical tests were applied to this summary visualization. D. Differential expression of MHC-II genes across diets and reversal conditions. Points show gene-level log2 fold changes for H2-Aa, H2-Ab1, and Cd74, and gray bars indicate the mean log2 fold change across these genes within each cell type and condition. Differential expression was computed separately for each cell type and condition versus control using Libra single-cell Wilcoxon testing with mouse as the replicate variable and Benjamini-Hochberg multiple-testing correction. The analysis included male cells only (n = 68,171 cells from 23 mice total across all conditions shown); each comparison used n = 3 control mice and n = 2 mice in the corresponding diet or reversal group. No additional statistical tests were applied to this summary visualization.

We observed even stronger associations between MHC-II genes and diet in the Wilcoxon DEGs (Fig 7d). Of particular note, MHC-II gene (H2-Aa, H2-Ab1, and Cd74) expression appeared to be repressed (FDR < 0.05) even following both the four and nine month dietary reversals, across diverse cell types. A subset of these diet-gene-celltype associations were additionally FDR-significant and/or trending similarly in the DESeq2 analysis. This further indicates potential persistent metabolic reprogramming due to high-fat diets on immune status, which could have implications for long-term impacts of dietary changes on host health.

## Discussion

In this study, we present a high-resolution, multi-omic atlas of host-microbe interactions, constructed by subjecting mice to seven compositionally diverse high-fat diets and a purified low-fat control diet. This work maps microbial and host responses across two longitudinal cohorts with discrete baseline microbiomes over the course of a year. This dual-cohort design allows us to disentangle how the initial ecosystem structure dictates the trajectory of diet-induced microbiome remodeling. Furthermore, by reverting a subset of animals to control diet, we decoupled transient adaptations from persistent biological signatures. Integrating longitudinal metagenomics, metabolomics, lipidomics, and host single-cell transcriptomics, this resource provides the community with a robust reference for how specific dietary lipids remodel the gut ecosystem and host physiology across time and varying microbial contexts.

This resource extended prior efforts in understanding the associations between host sex, diet, time, and the gut microbiome^10,53–55^. Across seven high-fat diets, fat composition emerged as the dominant driver of microbial structure, exceeding sex and aging effects at the species level, consistent with prior demonstrations that dietary fat source reproducibly reshapes gut community ecology^8,56^. Although several diets transiently increased alpha diversity, species-level modeling revealed that high-fat feeding more consistently depleted shared commensals than it generated conserved enrichments, and longitudinal clustering showed that early enrichments often resolved into persistent losses. Approximately half of diet-associated taxa failed to fully recover after dietary reversal, echoing evidence that diet-induced microbial shifts can establish durable ecological states that are not readily reversed by nutritional normalization^26,57^. Hierarchical clustering further revealed three broad diet groupings, with coconut oil, palm oil, and lard clustering together; olive oil, milkfat, and fish oil forming a second cluster; and the ketogenic diet remaining distinct, underscoring both shared ecological constraints and fat-specific selection pressures. Importantly, baseline microbiome composition acted as a decisive filter for dietary response: fewer than 12% of species-level associations were conserved across *Helicobacter*-negative and *Helicobacter*-positive cohorts, and many taxa exhibited directionally discordant responses, supporting the concept that host–diet interactions are conditioned by pre-existing microbial networks^58,59^. Together, these findings indicate that high-fat diets impose progressive and partially persistent restructuring of the gut microbiome, but the magnitude and direction of these effects depend strongly on both fat composition and baseline community structure.

The fecal metabolite data revealed expected associations between specific fats and metabolic features; for example, the ketogenic diet was associated with increased ceramides, sphingomyelins, and acyl-carnitines consistent with enhanced fatty-acid oxidation^60^, whereas fish oil enriched EPA- and DPA-derived omega-3 metabolites and tocopherols^61^, and saturated-fat–rich diets preferentially increased palmitoyl-, stearoyl-, and oleoyl-linked lipid species and ceramides. Second, we observed numerous microbiome associations with diets as well as discrete metabolites with known ramifications for host health, indicating the potential in identifying microbial mediators of response to nutrition. *Alistipes spp.*, specifically *Alistipes finegoldii,* was increased across multiple diets; *notably, A. finegoldii* emerged as a central hub in microbe–metabolite association networks and was linked to extensive alterations in lipid, bile acid, polyphenol, and neurotransmitter metabolism, including carboxyethyl-GABA. In contrast, *Lactobacillus johnsonii,* another microbe considered to be protective against carcinogenesis^62^, was depleted across most diets (most prominently in the ketogenic diet), and its loss coincided with depletion of lactobacilli-associated metabolites such as lactobacillic acid and lactose derivatives. Additional microbes that were altered included *Akkermansia muciniphila* (most strikingly increased in the ketogenic diet). Other organisms with reported host health impacts that were impacted to varying degrees were *Bacteroides thetaiotaomicron* (increased; coconut oil [significant], generally increased across most diets except milkfat), *Bifidobacterium pseudolongum subsp. globulosum* (decreased; all diets except ketogenic), *Limosilactobacillus reuteri* (decreased; most or all diets), *Dubosiella newyorkensis* (decreased; all diets), and *Parasutterella excrementihominis* (decreased; most or all diets)^32,35,63–68^. Across diets, the dominant transcriptional shifts localized to the intestinal epithelium, particularly mature proximal enterocytes, and were characterized by coordinated changes in nutrient transport and metabolic genes such as *Slc2a2*, *Sis*, and *Cyp4a10*, together with enterocyte-associated lipid-handling genes including *Fabp1* and *Hmgcs2*. In parallel, epithelial defense and antigen-presentation genes, including *Reg3b*, *Cd74*, and *H2-Aa*, were altered across cell types, along with immune-modulatory factors such as *Ccl5* and *Rorc*, indicating that long-term dietary fat composition reshapes both metabolic and host–immune interface programs in the gut epithelium.

One of the core motivations for building this resource was to identify the degree to which dietary choices could have persistent effects on the host even following reversion to a control diet for a period of three or eight months, approximately equal to seven and 21 human years, respectively^69^. Our reversal analysis indicates that persistent changes may in fact occur, consistent with previous studies demonstrating microbiome “scarring” from antibiotic usage^70,71^ or persistent microbiome alterations that are associated with post-dieting weight gain^26^. While body weight and systemic metabolites mostly resolved upon return to a control diet, we observed that approximately 50% of diet-induced species failed to revert to baseline, with organisms like *Alistipes finegoldii* remaining persistently enriched and commensals like *Lactobacillus johnsonii* remaining persistently lost.

Notably, this ecological memory was mirrored in the host transcriptome, where single-cell RNA sequencing revealed sustained suppression of MHC-II pathway genes (e.g., H2-Aa, Cd74) in intestinal epithelial cells. Given that we and others have previously shown that epithelial MHC-II expression plays crucial roles in coordinating immune surveillance against tumors, enteric pathogens, and local inflammation, this sustained suppression implies a lasting state of vulnerability^20,72,73^. Collectively, these findings indicate dietary history as a distinct biological variable, where past nutritional stress creates a latent susceptibility mediated by both microbial and host alterations that persist despite lifestyle correction. Further work is needed to interrogate the causal direction of these persistent host-microbiome states, as well as their stability, in human models and in light of other exposures.

Overall, this resource will be valuable for researchers interested in identifying links between the microbiome, fecal and serum metabolome, lipidome, and host transcriptional state. We anticipate the associations and datasets here can be integrated to evaluate hypotheses that could not be easily tested in humans or in other observational cohort data. Our full resource is available at https://btierneyshiny.shinyapps.io/high-fat-diets-multi-omics/.

### Limitations of the study

There are several limitations of our study. First, some of the associations that are consistent across all high-fat diets could be confounded by the ingredients of the purified control diet, which is soybean oil based. Similarly, the ketogenic diet differs from the isocaloric high-fat panel in both caloric density and macronutrient composition. Second, as reported in prior high-fat diet microbiome studies, Lactococcus species can represent potential contaminants in some datasets^8^. In our study, the increase in lactococcal taxa was not uniform across diets, arguing against a simple global contamination artifact; however, this possibility should still be considered when interpreting taxon-specific findings. Third, the reversal experiments demonstrate persistence of selected phenotypes within the defined observation window, but they do not establish whether these effects remain stable over longer timescales. A further limitation is uneven sample depth across modalities. In particular, intestinal single-cell RNA-seq were available for fewer samples, which reduces power for direct cross-modality association analyses, especially microbe-to-host DEG relationships. These integrative analyses should therefore be interpreted as prioritization tools rather than definitive evidence of host–microbe coupling. Finally, this study was designed as a systems-level observational resource rather than a mechanistic dissection. The atlas therefore does not by itself establish causal relationships between dietary fat source, microbial community structure, metabolite profiles, and host tissue responses. Instead, its principal value lies in providing a comparative framework that can be used to generate testable hypotheses and guide focused causal experiments.

---

**Supplementary Figure 1.**
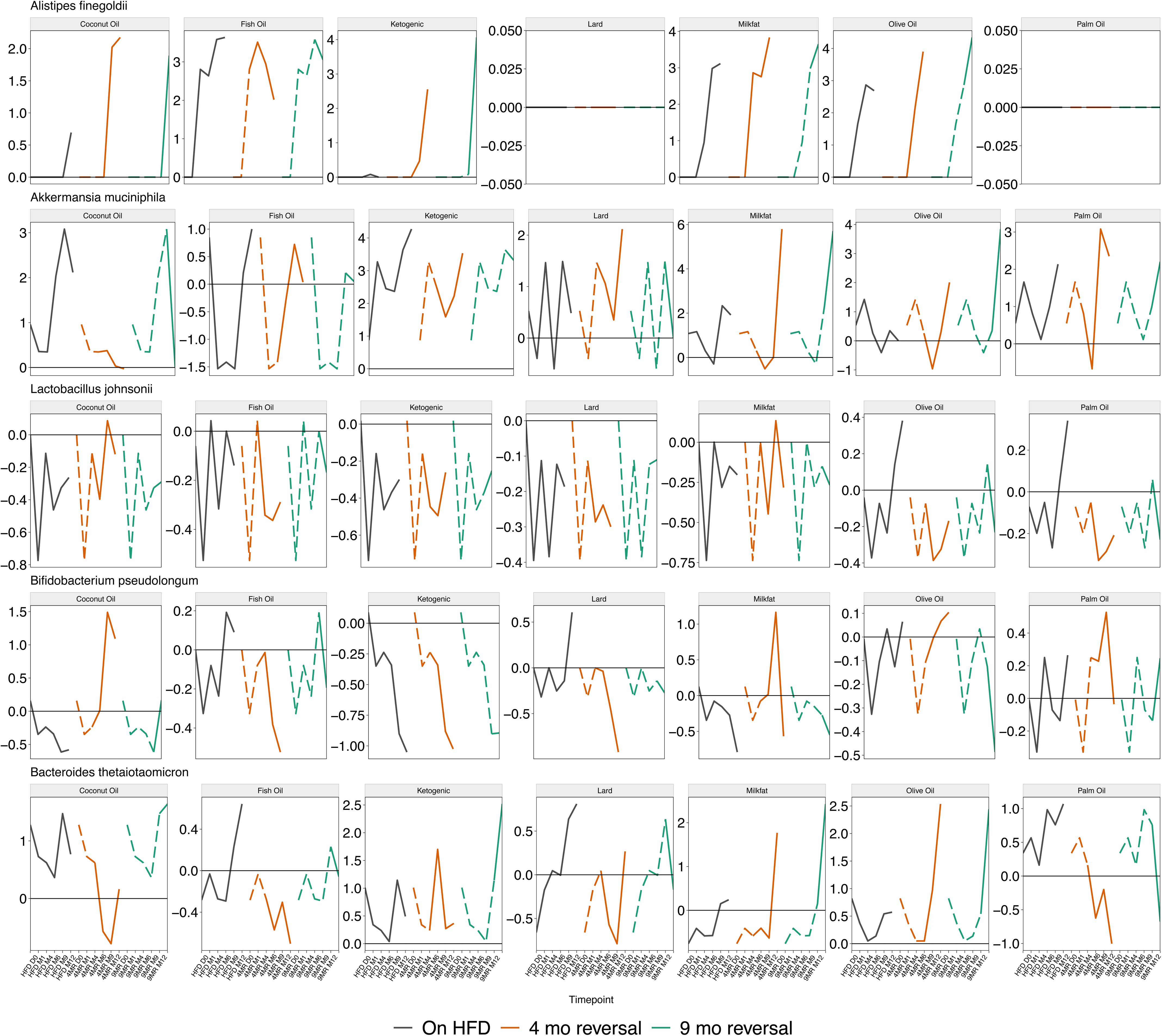
Example trajectories for taxa with persistent reversal or depletion patterns (Cohort 2, H−; n = 158 mice, 809 fecal samples; arms as in Fig. 3). Each mini-panel is one diet–taxon combination. Left segment, average log2 fold change versus time-matched control on continuous high-fat diet (non-reversal). Middle, four-month reversal (dotted, on high-fat before switch; solid, after return to control). Right, nine-month reversal (same convention). Black, continuous high-fat; orange, four-month reversal cohort; green, nine-month reversal cohort.

**Supplementary Figure 2.**
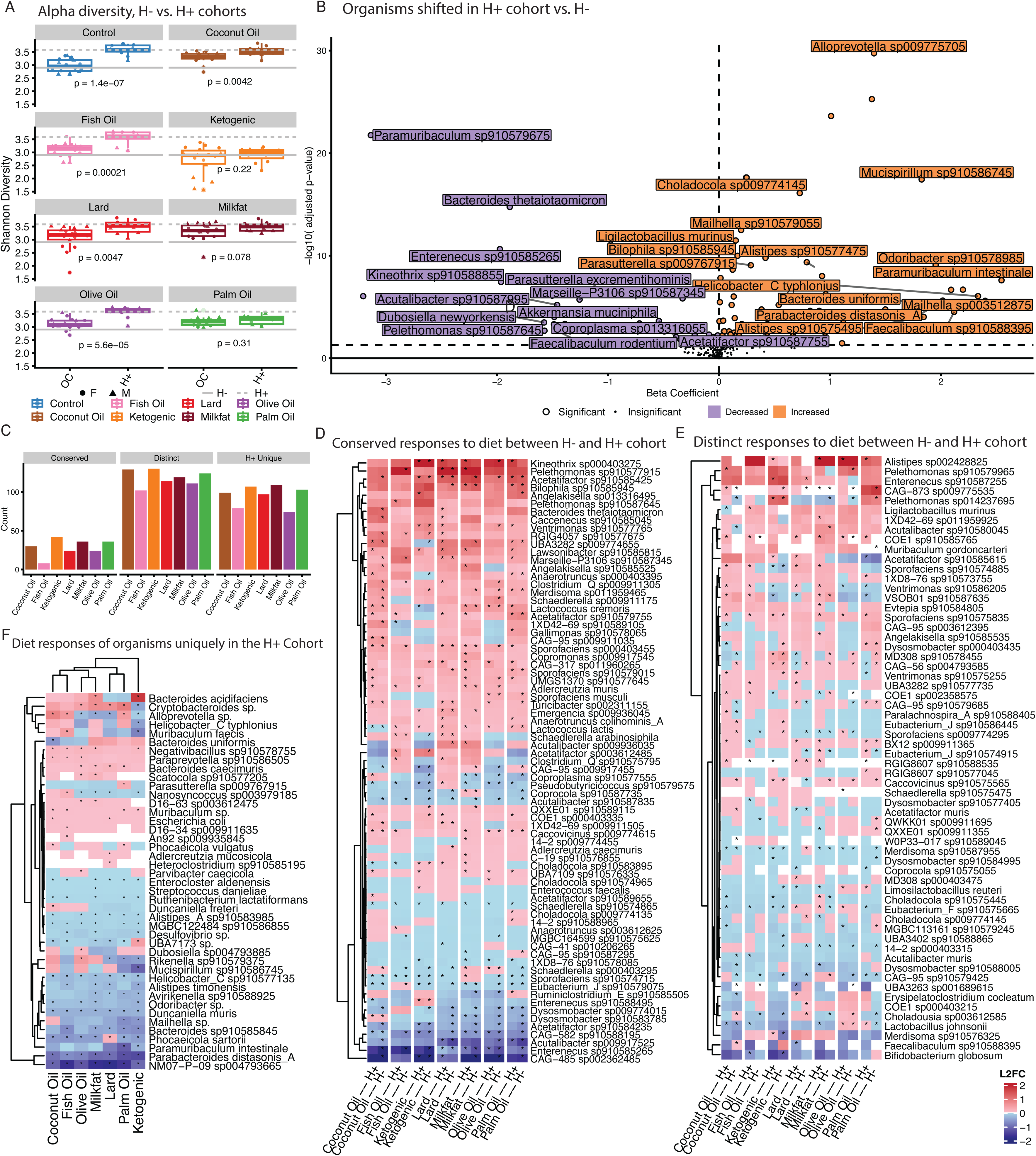
In-house (H+) versus Jackson (H−) cohort comparison using longitudinal fecal microbiome data (H−: n = 158 mice, 809 samples; H+: n = 64 mice, 302 samples). A. Alpha diversity by cohort and diet (H− reference line solid, H+ dashed). Unadjusted p-values from t-tests in-panel. Diet colors match panel C. Point shape, sex (M/F) (all samples per cohort as above). B. Volcano plot for H+ versus H− (control-diet fecal samples only): per-species linear mixed model ABUNDANCE ∼ COHORT + (1|ANIMAL_ID), Benjamini–Hochberg across species; dashed line, FDR 0.05 on −log10 scale. X axis, cohort coefficient; Y axis, −log10(adjusted p-value); color, direction of significant hits. C. Conserved, distinct, and unique diet–microbe association counts across cohorts (colors as panel A). Asterisks elsewhere in this figure, BH FDR < 0.05. D.-F. Heatmaps of log2 fold change versus H− reference for the three panel C categories. X axis, diet–cohort combinations; columns grouped by diet. Asterisks, BH FDR < 0.05 in that cohort. Red/blue, L2FC versus control within cohort. Panel D, same-direction L2FC across cohorts; panel E, at least one opposing L2FC between cohorts.

**Supplementary Figure 3.**
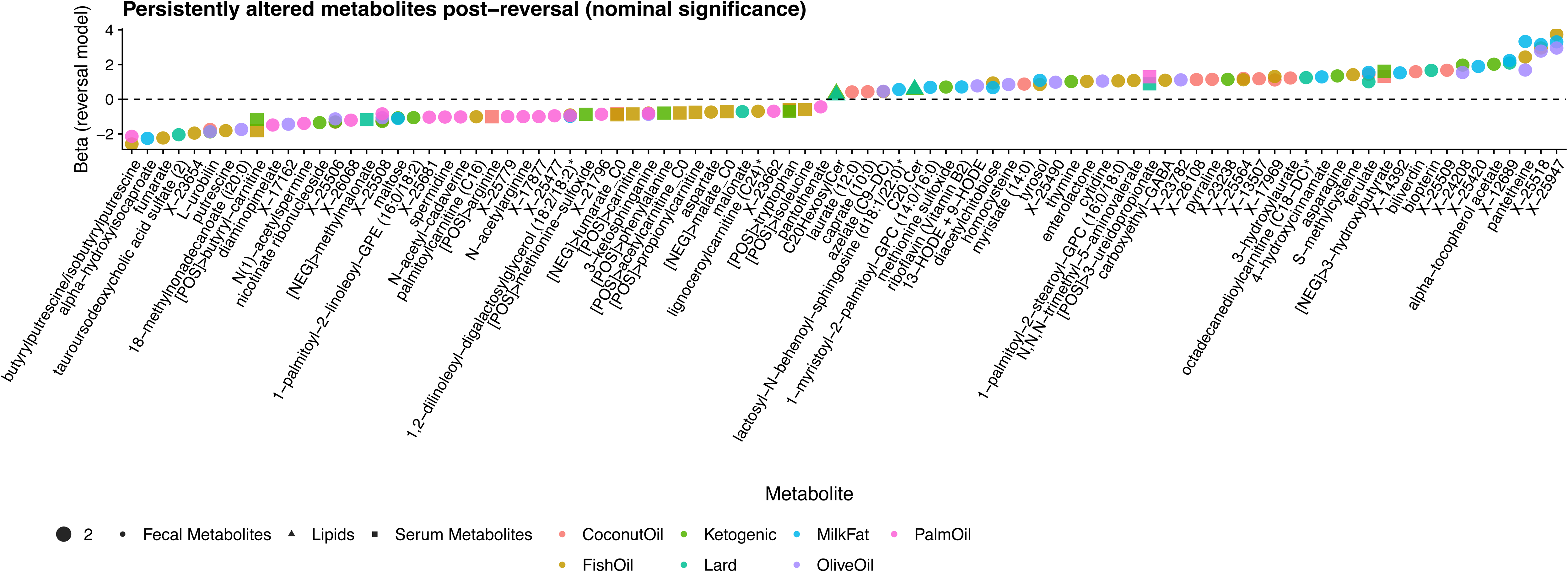
Metabolites altered after reversal at nominal p < 0.05 (extends Fig. 4, which used FDR). Each point is one feature; X axis, feature; Y axis, post-reversal association strength; color, diet; shape, layer (fecal metabolite, lipid, serum). Horizontal dashed line, zero.

**Supplementary Figure 4.**
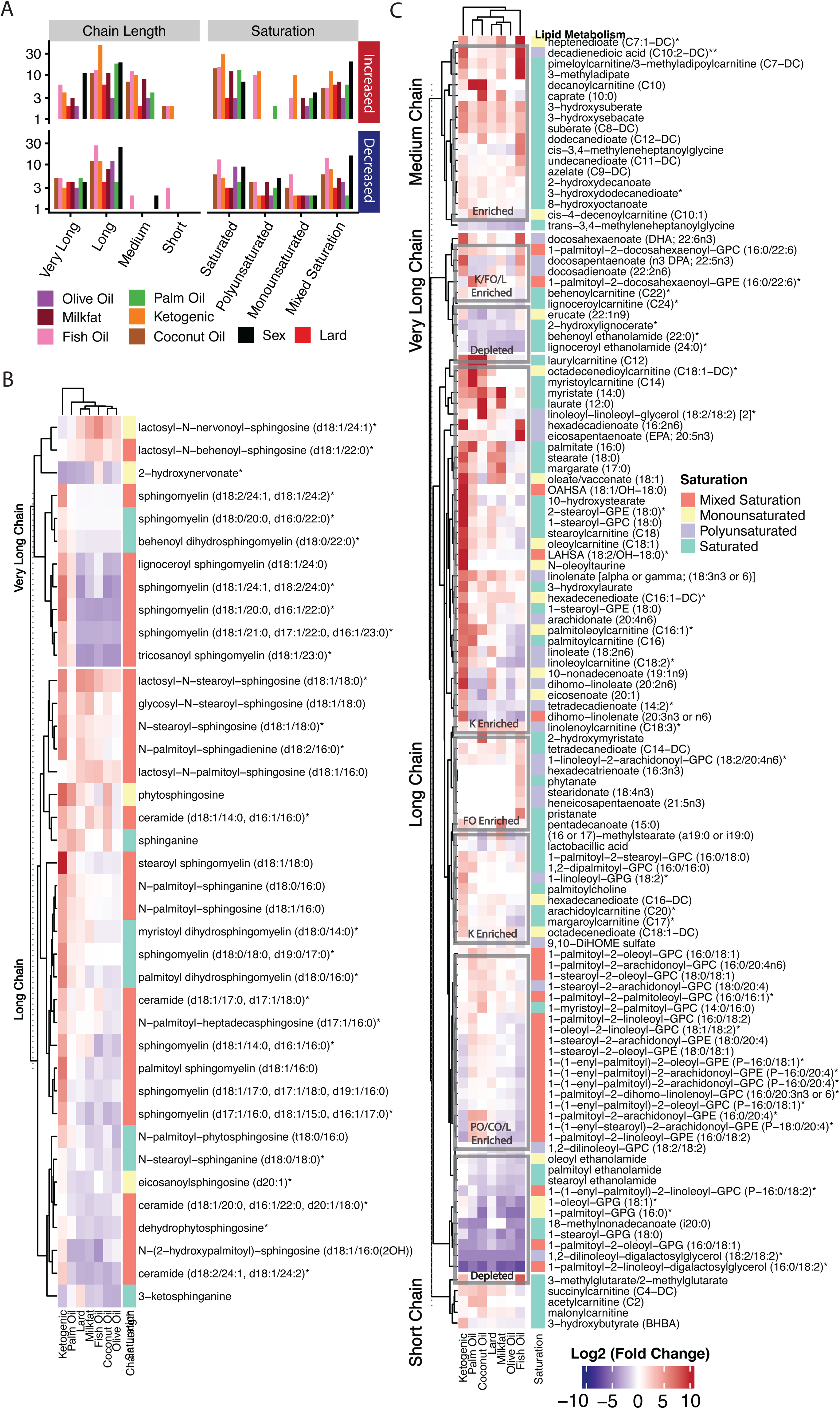
Additional fecal lipid detail (same modeling framework as main lipid panels). A. Lipid feature counts by chain length and saturation class, split by association direction (red, increased with term; blue, decreased). Bar height, count; Y axis log-scaled. Fill, model term (diet or sex). B. Sphingolipids with Benjamini–Hochberg FDR < 0.1 for at least one contrast (rows clustered; columns as panel C: diets then sex). Cell color, mean L2FC; asterisks per plotting rules on the figure. C. Non-sphingolipid lipids meeting the same FDR rule; layout as panel B; saturation strip beside rows; grey boxes, manual interpretive highlights only.

**Supplementary Figure 5.**
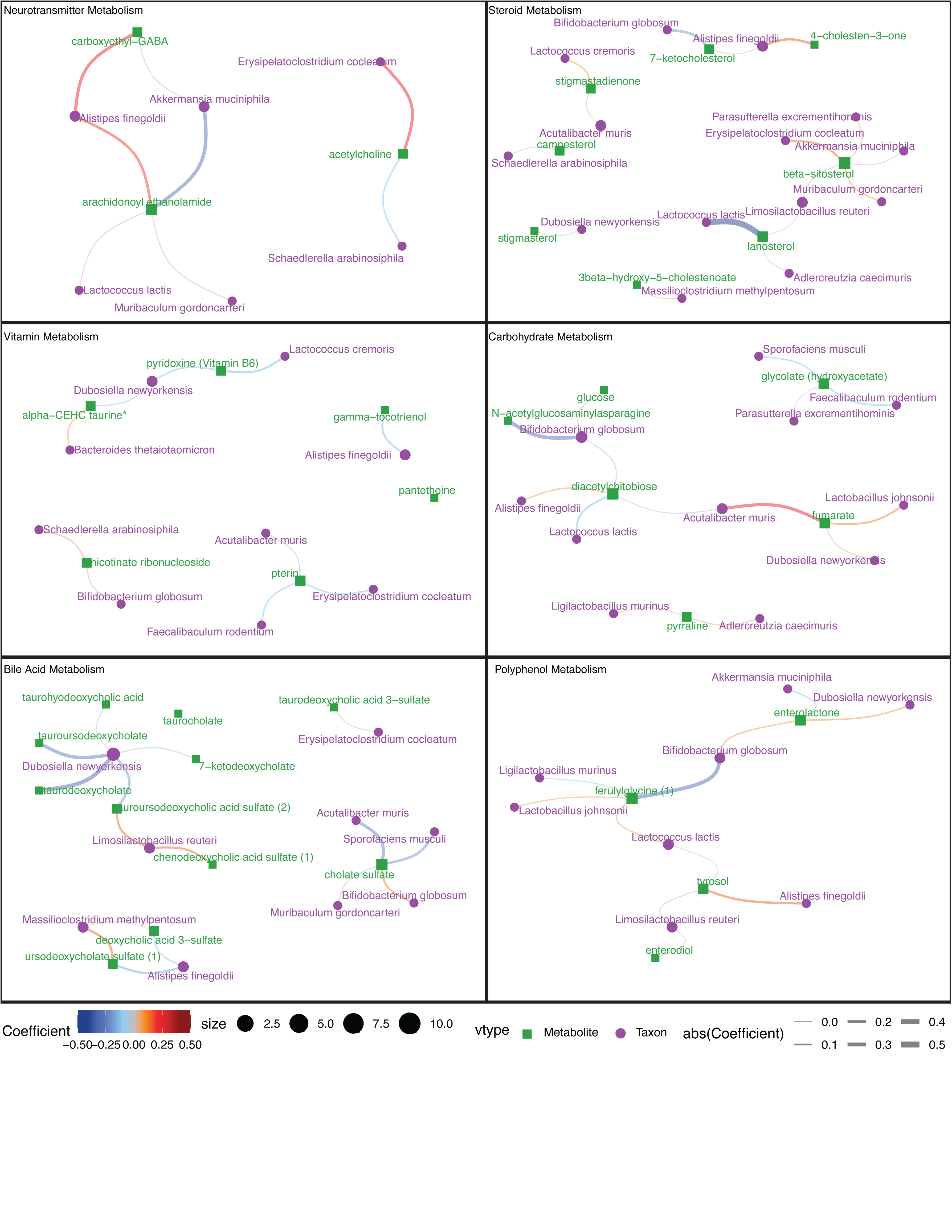
Microbe–metabolite networks by fecal metabolite class (as Fig. 6: |LASSO coef| > 0.1). Red/blue, sign of association; width, |coef|; green squares, metabolites; purple circles, taxa; size, degree.

**Supplementary Figure 6.**
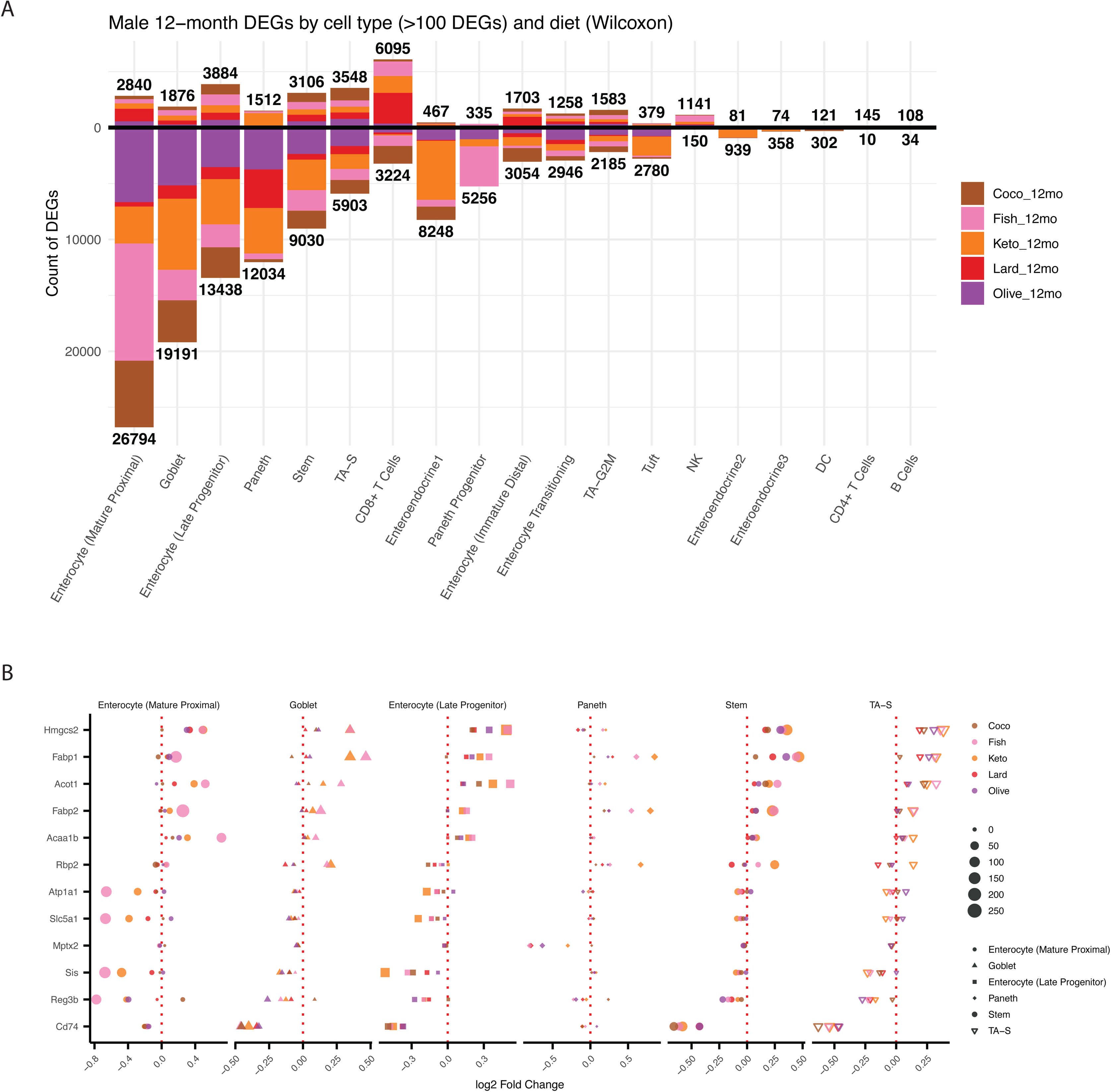
Wilcoxon-based single-cell differential expression (Libra), complementing Fig. 7. A. DEG counts by cell type and diet (12-month males; FDR < 0.05; mouse as replicate; Benjamini–Hochberg) (n = 3 control and n = 2 per diet group; 32,895 cells across listed diets; cell types with <50 cells in a comparison excluded). No further tests on this summary plot. B. Top five Wilcoxon DEGs in the six cell types with the most hits (ranked by |log2FC| among FDR-significant genes). Points, log2FC; size, −log10(FDR); color, diet. Same testing framework as panel A (n = 3 control, n = 2 per diet per comparison). No further tests on this summary plot.

## Data and Code Availability

The 10X-formatted fastq files for single-cell RNA-sequencing have been deposited into the European Nucleotide Archive (accession number: PRJEB105080). The fecal metagenomics raw data has also been deposited into the European Nucleotide Archive (accession number: PRJEB106128). Metadata sheets associated with these submissions are available at: https://github.com/b-tierney/multi-omics-of-high-fat-diets/. This repository additionally contains scripts for analysis of the processed data. Full taxonomic, metabolomic, and functional data are available at https://figshare.com/projects/_b_Diverse_high-fat_diets_drive_multi-omic_reprogramming_that_persists_after_dietary_reversal_b_/270115 as well as https://btierneyshiny.shinyapps.io/high-fat-diets-multi-omics/.

## Author Contributions

Conceptualization, S.B.; Data curation, B.T.T., S.B., C.Meydan., and A.G.V.C.; Investigation: S.B., O.E., K.A.O, K.P., S.S., H.A., I.E., C.C., V.S., B.Y., M.R.F., C.D., C.Mozsary, E.K., A.G., N.D, D.N., K.A.R., D.J.B..; Formal analysis, B.T.T., S.B., C.Meydan., A.G.V.C., and E.O.; Statistical design, B.T.T., S.B., C.Meydan., A.G.V.C., and E.O.; Software, B.T.T., S.B., C. Meydan., and A.G.V.C.; Visualization, B.T.T., S.B., C.Meydan., A.G.V.C., E.O., and J.P.; Supervision, B.T.T., S.B., and C. Meydan, C.E.M.; Resources: C.A.T, C.J.P.; Project administration, B.T.T., S.B., C.Meydan., C.E.M., and T.M.N.; Funding acquisition: S.B., K.B., C.E.M, B.T.; Writing – original draft, B.T.T., S.B., C.Meydan., and A.G.V.C.; Writing – review and editing, all authors.

## Acknowledgements

We thank Cold Spring Harbor Laboratory Cancer Center Shared Resources (Animal, Single Cell Sequencing and Sequencing Core Facilities) supported in part by the National Cancer Institute Cancer Center Support Grant 5P30CA045508. S.B was supported by the National Cancer Institute (R37CA292807), STARR Cancer Consortium (#I13-0052), Oliver S. and Jennie R. Donaldson Charitable Trust, the G. Harold and Leila Y. Mathers Charitable Foundation, the Mark Foundation for Cancer Research (20-028-EDV), Chan Zuckerberg Initiative/Silicon Valley Community Foundation (2021-239862), the Cold Spring Harbor Laboratory and Northwell Health Affiliation, Swim Across America. A.G.V.C. was supported by 5T32HG002295. T.M.N. was supported by a Medical Scientist Training Program grant from the National Institute of General Medical Sciences of the National Institutes of Health under award number: T32GM152349 to the Weill Cornell/Rockefeller/Sloan Kettering Tri-Institutional MD-PhD Program. The computations by T.M.N. in this paper were run with HPC resources supported by the Scientific Computing Unit at Weill Cornell Medicine. B.T.T. was supported by the Advanced Research Projects Agency for Health (ARPA-H) under award number D24AC00345-00. ARPA-H provided 90% of total costs with an award total of up to $18,474,445. NIEHS R01 R01ES032470 and NIDDK R01DK137993 provided 10% of support. The content is solely the responsibility of the authors and does not necessarily represent the official views of the Advanced Research Projects Agency for Health or the National Institutes of Health. The lipidomics analyses were performed at the Medical University of South Carolina’s Lipidomics Shared Resource supported by NIH (C06 RR015455), Hollings Cancer Center Support Grant (P30 CA138313), or Center of Biomedical Research Excellence (Cobre) in Lipidomics and Pathobiology (P30 GM103339). The metagenomic sequencing was completed at the Weill Cornell Medicine Genomics Core Resources Facility.

## STAR Methods

### Mice, diets and study design

C57BL/6J (strain name: 000664) mice were obtained from the Jackson Laboratory. Mice were housed in pathogen-free conditions and maintained at 12 hours light/dark cycles. All mice in this study were age- and sex-matched (both male and female). Food and water provided ad libitum. All animals used in this study were handled according to ethical procedures approved by The Institutional Care and Use Committee (IACUC) at Cold Spring Harbor Laboratory.

### Diet design

We have designed custom isocaloric HFDs (Envigo, 4.6 kcal/g total, 22.7% fat by weight, 45% kcal from fat) that have identical macronutrient (protein, carbohydrate and fat) and micronutrient (vitamins and minerals) composition with different fat sources and a low-fat purified control diet (Envigo, 3.8 kcal/g, 7.2% fat by weight and 17% kcal from fat, soybean oil based). In addition, to study how fat abundance in diet affect outcomes, we designed a lard-based ketogenic diet with very high fat and low carbohydrate abundance (Envigo, 7.0 kcal/g, 70.2% fat by weight, 90.3% kcal from fat and fat to protein and carbohydrate ratio is approximately 4:1). The detailed composition of each diet can be found in Supplementary Table 1.

### Study design

Experiments were performed using age- and sex-matched C57BL/6J mice starting at 8 weeks old housed in specific pathogen-free (SPF) conditions. To assess the impact of baseline microbiome composition on dietary responses, we utilized two distinct microbial cohorts:

- **Helicobacter-negative (H-) Cohort:** C57BL/6J mice were purchased from The Jackson Laboratory. These mice are standard SPF flora-competent but historically devoid of *Helicobacter* species and other pathobionts.
- **Helicobacter-positive (H+) Cohort:** We utilized an in-house breeding colony of C57BL/6J mice maintained at Cold Spring Harbor Laboratory. Unlike the H− cohort, these mice naturally harbor a distinct commensal community including *Helicobacter* species. PCR and metagenomic analysis of fecal contents confirmed in this cohort the presence of *H. typhlonius* and *H. mastomyrinus*, species known to associate with elevated epithelial MHC-II expression^20^.

To prevent cross-contamination of microbiomes, H− and H+ cohorts were housed in separate, isolated containment rooms for the duration of the study.

We employed a longitudinal study design spanning 12 months to assess both the chronic impacts of high-fat feeding and the persistence of these signatures after dietary cessation. Before starting high-fat diets, all mice were placed on a purified control diet for two weeks. The study was divided into three arms:

1. **Continuous Feeding Arm:** Mice were maintained on their respective HFD or control diets for the full duration (up to 12 months) to establish the chronic trajectory of host-microbiome interactions.
2. **Early Reversal Arm:** After 4 months of HFD feeding, a sub-cohort of mice was switched to the Low-Fat Control diet to assess phenotype reversibility.
3. **Late Reversal Arm:** After 9 months of HFD feeding, a second sub-cohort was switched to the Low-Fat Control diet to assess phenotype reversibility.

### Metabolic cage measurements

Age-matched C57Bl/6 mice were stratified by diet and sex to ensure nearly equal numbers of each per diet. Mice were placed into metabolic cages accommodating one mouse each (CLAMS, Columbus). While in metabolic cages, original diets for mice were continued and mice were habituated to the cages for one week before testing under 12h each light/dark cycle. Within metabolic cages, acquired parameters included O_2_ volume, O_2_ in/out, accumulated O_2_, CO_2_ volume, CO_2_ in/out, accumulated CO2, RER, Heat, Flow, Pressure, Feeder weight, feeder accumulation, drink weight, drink accumulation, movement in X axis, movement in Y axis and movement in Z axis. Mice were kept within metabolic cages for 4 days in each time interval (1m, 3m, 7m, 12m) and data was recorded every 5 minutes for each parameter.

### Metabolic cage bioinformatics analysis

Metabolic cage data was recorded in CLAMS format by Columbus Instruments and converted to CSV files using the OXYMAX software. Samples that experienced technical issues in data recording and/or animal death were removed from the dataset. For the analysis and visualizations, we used R version 4.2 along with the following packages: dplyr, tidyr, ggplot2, tidyverse, ggrepel, RColorBrewer, ComplexHeatmap, and circlize. Data was filtered using the RER parameter to be within the biologically relevant range of 0.5 to 1. Samples were grouped by diet, sex, and time intervals, and the average RER values were calculated over all measurements.

For activity analysis, to ensure consistent measurements across samples, the interval was filtered to at most 1000 measurements. Groups were formed by sex, diet, night/day cycle, and time. Total ambulatory activity was calculated by taking the average of the X.ambulatory and Y.ambulatory parameters.

### SnapFreeze RNA-isolation and bulk RNA sequencing

Mice were euthanized within a CO2 chamber and the small intestine was removed and separated from intraperitoneal fat. A cold (4 °C) wash with 1xPBS removed debris and feces. A small portion of the ileum was cut and snap-frozen in liquid nitrogen. RNA was isolated using a RNeasy Plus Mini Kit (Qiagen 74134) and gDNA was removed using the provided elimination column.

Total RNA samples underwent quality control and normalization prior to library construction. Host ribosomal RNA was depleted using the NEBNext rRNA Depletion Kit (Human/Mouse/Rat; New England Biolabs, E6310). Strand-specific libraries were prepared with the NEBNext Ultra II Directional RNA Library Prep Kit for Illumina (New England Biolabs, E7760) according to the manufacturer’s protocol. Library fragment size distributions were assessed on an Agilent TapeStation using the High Sensitivity D1000 ScreenTape assay (Agilent Technologies, 5067-5584 and 5067-5585). Libraries were pooled and sequenced as paired-end 150 bp reads (PE150) on an S4 flow cell using the Illumina NovaSeq 6000 sequencing platform at the Genomics Resources Core Facility (GRCF) at Weill Cornell Medicine.

### Single-cell sequencing

Isolation of intestinal crypts was conducted following previously established protocols. In short, the entire intestine was excised, with fat, connective tissues, and blood vessels removed, and flushed with ice-cold 1X PBS. After orienting the tissue, the small intestine was sectioned into 3-5 cm fragments and incubated in 1X PBS/EDTA (7.5 mM) with gentle agitation at 4°C for 30 minutes. Crypts were released from the tissue by mechanical disruption, passed through a 70-micron strainer to eliminate villi and tissue debris. The crypts were then washed with cold PBS and spun down at 300g for 5 minutes. To obtain intestinal epithelial cells (IECs), the crypts were dissociated into single cells using TrypLE Express (Cat# 12604-013, Invitrogen). Dead cells were excluded from analysis using the viability dye SYTOX (Cat# S34857, Life Technologies), and the SYTOX-negative population was sorted using a Sony SH800S cell sorter. single-cell droplets were immediately prepared on the 10X Chromium according to manufacturer instructions at Cold Spring Harbor Laboratory Single Cell Facility. Single cell libraries were prepared using a 10X Genomics Chromium Controller (Cat #120223, 10X Genomics) and the 10X Genomics Chromium Next GEM Single Cell 3’ Gene Expression kit (Cat #1000268, 10X Genomics) according to the manufacturer’s instructions.

### Fecal metagenomics

Following collection of fecal samples, samples were snap frozen in liquid nitrogen and stored at −80 °C. DNA extraction was carried out using a fecal sample-based modification of the Maxwell® RSC PureFood GMO and Authentication Kit (Catalog No. AS1600; Promega Corporation, Madison, WI) following the manufacturer’s guidelines. In brief, 75 mg of fecal pellets were placed into 2 mL microcentrifuge tubes, to which 1 mL of CTAB buffer was added, followed by vortexing with the MO BIO Vortex Adapter for 30 seconds. After homogenization, the samples were heated at 95°C for 5 minutes, cooled at room temperature for 2 minutes, and vortexed horizontally for an additional minute. The pellets were then manually homogenized in the same tubes using disposable Fisherbrand™ Pellet Pestles (Thermo Fisher Scientific, Waltham, MA, USA) until fully dispersed. For further lysis, 40 μL of Proteinase K and 20 μL of RNase A were added, and the samples were vortexed for 1 minute before incubating at 70°C for 10 minutes. Extracted DNA was used for shotgun metagenomic library preparation using standard Illumina DNA Prep chemistry, and sequenced on an Illumina NovaSeq platform.

### Gut microbiome bioinformatic analysis

Paired-end stool microbiome samples were quality-controlled with a pipeline wrapping portions of the bbtools suite (version 38.92) that we have leveraged in other work^74,75^. Initially, clumpify [parameters: optical=f, dupesubs=2, dedupe=t] grouped the reads. Subsequently, bbduk [parameters: qout=33, trd=t, hdist=1, k=27, ktrim=“r”, mink=8, overwrite=true, trimq=10, qtrim=’rl’, threads=10, minlength=51, maxns=-1, minbasefrequency=0.05, ecco=f] was used to excise adapter contamination. Tadpole [parameters: mode=correct, ecc=t, ecco=t] was used to correct sequencing errors. The repair function of bbtools was used to remove any unmatching reads. Finally, to eliminate potential human contaminants, Bowtie2 [parameters: --very-sensitive-local] was employed for alignment to the human genome.

To generate alignment-based taxonomic annotations for our database, we used three different approaches, each with the default settings: MetaPhlAn4 (V4.0.4, aligned against the default database), Kraken2/Bracken (V2.1.2, aligned against against the complete genomes for microorganisms on RefSeq as of October 2022), and Xtree (V0.92i, aligned against the Genome Taxonomy Database r207).^76–80^ Xtree, a k-mer based aligner, produces coverage statistics, including global coverage (total genome coverage) and unique coverage (reads mapping to a specific genome in the database). Bacterial genomes were filtered based on a threshold of 10% global and/or 5% unique coverage. The relative abundances for Xtree were calculated by dividing the total reads aligned to a specific genome by the total number of reads aligning to all genomes. MetaPhlAn4 automatically computed relative abundances. For the Kraken2 output, Bracken was utilized to determine relative abundances. These classifiers each use different databases and provide slightly different perspectives on the composition of the samples in question so, as a result, we opt to provide all three of their outputs in the resource described here. In the main text, however, we report results from MetaPhlAn4, as it is well-established for gut microbiome studies and, based on this study and our prior work^74^, tends to have a reasonable tradeoff between sensitivity and specificity. Alpha and beta diversities were computed using the vegan package in R on log-transformed relative abundance data ^81^. Unless otherwise noted, software was executed with default settings.

### LC-MS analysis of polar plasma metabolites

Plasma samples (10 µl) were extracted with 90 µl of cold acetonitrile: methanol: formic acid (75:25:0.2, v/v/v) containing ^15^N- and ^13^C-labeled amino acid standards (MSK-A2-1.2, Cambridge Isotope Laboratories, Inc). Samples were vortexed vigorously followed by centrifugation for 30 minutes (20,000 x g, 4°C). The supernatant was transferred to an LC vial and 5 µl was injected onto a ZIC-pHILIC (150 × 2.1mm, 5 μm particle size; EMD Millipore) column. Polar extracts were separated on a Vanquish UHPLC coupled to a QExactive orbitrap mass spectrometer as previously described^82^. The samples were analyzed in a randomized order with QC samples injected every 4 hours for batch normalization. A pool of all the biological samples was prepared and analyzed using a data-dependent acquisition method. The raw data were searched against an internal metabolite library containing >200 polar metabolites using a 2-ppm mass tolerance and a 6 second retention time threshold relative to authentic standards. Relative metabolite abundances were quantified using Skyline Daily (v21.1).

### Lipidomics analysis

Lipidomics analysis were performed by LC-MS/MS followed the protocol described previously^83^. Briefly, samples were supplemented with internal standards, and 2mL of <u>isopropyl alcohol</u>: water: <u>ethyl acetate</u> (30:10:60; v/v:v) was added to the <u>extracts</u>. Samples were subjected to two rounds of vortex and sonication followed by 10-min <u>centrifugation</u> at 4,000 rpm. The supernatant or top layer was used as lipid extract and subjected to LC-MS/MS for analysis of sphingolipid species. The MUSC <u>Lipidomics</u> Shared Resources performed lipid extraction and analyses. Inorganic phosphates (Pi) or total protein amounts were used for normalization.

### Fecal metabolomics (Metabolon)

Global untargeted metabolomics of mouse fecal samples was performed by Metabolon, Inc. (Durham, NC, USA) using ultrahigh-performance liquid chromatography–tandem mass spectrometry (UHPLC–MS/MS) platform. Frozen fecal samples were shipped on dry ice and stored at −80L°C until processing. Samples were prepared at a constant mass-to-volume ratio, and recovery standards were added prior to extraction for quality control. Proteins were precipitated with methanol under vigorous shaking to dissociate small molecules bound to protein matrices and recover chemically diverse metabolites. Following centrifugation, the resulting supernatant was divided into multiple fractions for analysis under positive and negative ion conditions across reverse-phase (C18) and hydrophilic interaction (HILIC) chromatographic columns.

### Microbiome-diet associations (non-reversal)

We used a linear modeling approach to identify putative associations between microbiome taxonomic relative abundances and dietary intake. In brief, for each timepoint and for each taxon identified in a given cohort, we fit the following model:

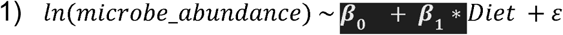

The *microbe_abundance* variable is the relative abundance of a given microorganism where all zero values have been replaced with a pseudocount of 0.5. The diet term corresponds to a categorical variable with 8 levels, one for each high-fat diet and another for the control. The control diet was the reference group. We adjusted for multiple testing across all models fit for a given taxonomic classifier using the Benjamini-Hochberg approach. An association was considered statistically significant at q < 0.05 and trending towards significance at q < 0.1. For the results highlighted in the manuscript, we reported taxa that were statistically significant or trending with a given diet and associated with at least two timepoints in the same direction.

While numerous modeling strategies could be employed to analyze this dataset (e.g., non-linear methods, mixed models incorporating multiple timepoints), we opted for a timepoint-by-timepoint linear approach due to 1) its simplicity in identifying timepoint-specific microbiome shifts, the ability to do statistical inference, and 3) and integrating the reversal analysis and non-reversal analysis into a single, straightforward pipeline. The major downside to this approach is a loss of power due to the number of associations fit and reliance on p-values, resulting in excessive conservatism in reporting results.

For the time and sex-associated counts in Figure 2D, we fit the following two additional models, extracting the number of FDR-significant associations for the sex and time effects (β_2_), respectively:

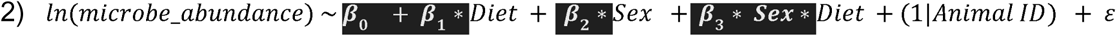

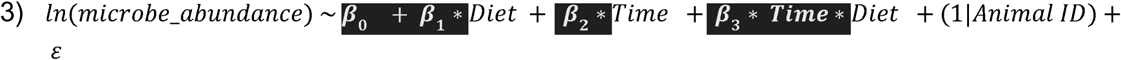

We opted to fit these separate models to consider sex and time to avoid using one, large overparameterized model with excess degrees of freedom and associated loss of power. Having 1) additionally enabled a straightforward way to compare the reversal versus non-reversal associations, matching timepoints in the different mice.

Analysis of the reversal cohorts was with the same modeling strategy to the non-reversal cohorts. We fit the same model specification, annotating for a given model, however, if a mouse was on or off the diet referenced in the categorical variable. For example, mice that underwent the four-month reversals would be considered “on diet” at month 1 and “off diet” at month 9. The interpretation of the beta coefficient on the diet variable, therefore, would enable insight into the impact of the diet after it was reverted to control. For example, if the association between the ketogenic diet and a certain organism was FDR significant and negative relative to control both before and after reversal, the interpretation would be that the impact of the ketogenic diet on that organism lingered. If an association were negative before reversal and non-significant, positive, or extremely low in absolute value, after reversal, that would indicate a reversion to normality.

### Longitudinal analysis of microbiome trends

Filtering for taxa highlighted as trending or significant by the statistical analysis, we used c-means clustering to identify temporal signatures of microbiome response to dietary changes in both the reversal and non-reversal cohorts. For the non-reversals, we selected microbial taxa that were associated with (q < 0.1) at least two on-diet timepoints. For the reversals, we selected taxa that were associated with at least two timepoints after the four month reversals and one timepoint (the only one available) after the nine month reversals. Before clustering, we computed the L2FC of each taxon relative to the control diet. These values are what are visualized in Figure 3. For the clustering itself (cmeans function in R, iter.max set to 2,000), we tested c-means for c = 16, c = 18, and c = 20. We report the values for c = 20 in the text, though the results were similar for all three in terms of exemplar trends displayed and the taxonomic/dietary compositions of each.

### Analysis of dietary alterations as a function of variation in baseline microbiome composition

We again used a linear modeling strategy to analyze the impact of differences in baseline microbiome on response to diet as described in the study design section above. We fit three models, using the same pseudocount replacement of 0.5 for 0 values as above, filtering for microbes that were present in more than three samples. In all models below the Cohort variable is a binary encoding of the BC versus the H+ the latter referring to mice with a distinct baseline microbiome due to their source lab. Animal_ID is a unique identifier for each mouse. We adjusted p-values for multiple hypothesis testing as above and used the same cutoffs.

For all timepoints and both cohorts, we fit a mixed model comparing the organisms abundant in each cohort.

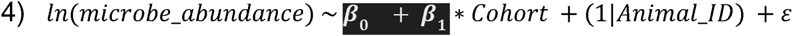

We next used data from just the H+ cohort in order to ascertain the within-cohort diet-specific effects:

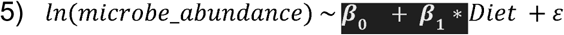

Finally, we used data from both the H+ cohort and original cohort to fit an interaction model to gauge diet-specific differences.

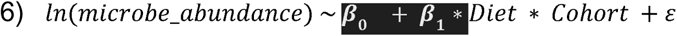

We next sought to categorize cohort-associated taxa into three groups that correspond to the three heatmaps in Supplementary Figure 2: 1) Same response to diet between cohorts, 2) Discrete response to diet between cohorts, 3) Organisms uniquely found in the H+ cohort. For group 1, we identified organisms/diets with q < 0.1 in equation 4 and with the same direction of beta coefficients between the cohorts. For group 2, we identified organisms/diets with q < 0.1 in equation 4 and opposite beta coefficients between the cohorts. For group 3, we identified organisms with q < .1 in equation 3) that were not found in the BC.

### Diet-microbiome-metabolite associations

We computed associations between diets and metabolites using another linear modeling approach of the following form:

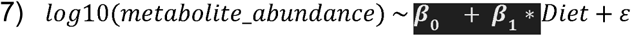

*Metabolite_abundance* is the mean of the metabolite abundance of a given fecal metabolite across the two timepoints metabolomics was completed (5 and 7 months). The *Diet* variable is defined categorically as in previous regressions. P-values and significance cutoffs were adjusted and defined as before, respectively.

We computed associations between microbial features and metabolites using two different methods: as LASSO regression and another linear modeling approach. For the LASSO regression, for each metabolite we used the cv.glmnet function from R’s glmnet package on centered and scaled microbe and metabolite, log10 transformed, relative abundance data. We replaced 0 values with a 0.5 pseudocount. We fit the following model, where *n* is the total number of microbes. To reduce the space of microorganisms we were interrogating, we only selected non single-genome-bin (SGB) organisms (i.e., prioritizing species with cultured representatives). For metabolites, to reduce auto-correlation as well as the entire search space, we clustered (hierarchically, cutpoint of 0.75) based on abundance and selected, when possible, metabolites with annotations as cluster representatives. In the event a cluster had multiple annotated metabolites, the one with the highest prevalence was selected.

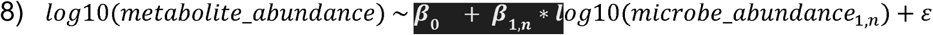

As a more conservative approach, reported in the Supplement, we fit the following linear model for each microbe-metabolite pair. Again, we used log10 transformations on the relative abundances of the microbes and metabolites, replacing 0 values with a 0.5 pseudocount and fit following model:

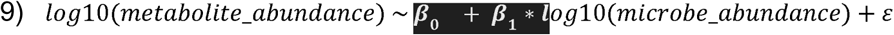

We defined significant/trending values and adjusted for multiple testing as before.

### Analysis of single-cell DEGs

Sequencing reads were aligned to the mouse reference genome and quantified using Cell Ranger (10x Genomics). The resulting count matrices were processed in R using the Seurat package. Cells were filtered based on the number of detected genes, total UMI counts, and mitochondrial gene content to remove low-quality cells and potential doublets. Gene expression data were log-normalized, and the top 2,000 highly variable genes were identified for downstream analysis. Cell cycle scores were computed using canonical S and G2/M phase markers, and cell cycle phase was assigned to each cell. To correct for batch effects across samples, datasets were integrated using Seurat’s canonical correlation analysis (CCA) workflow via FindIntegrationAnchors and IntegrateData. Principal component analysis (PCA) was performed on the integrated assay, and uniform manifold approximation and projection (UMAP) was used for two-dimensional visualization. Graph-based clustering was performed using the shared nearest neighbor (SNN) algorithm at a resolution of 0.9, yielding distinct clusters that were annotated into 21 cell types—including immune populations (B cells, NK cells, dendritic cells, plasmacytoid dendritic cells, CD4+ and CD8+ T cells), epithelial lineages (stem cells, transit-amplifying cells, enterocytes at multiple maturation stages, Paneth cells and progenitors, goblet cells, enteroendocrine subtypes, and tuft cells), and erythrocytes—based on known marker gene expression. The final dataset comprised 20,747 genes across 90,971 cells. Pseudobulk count matrices were constructed by summing raw reads across all cells for each mouse-cell type combination, and genes with less than 10 reads were filtered out. For each high-fat diet group with at least two male biological replicates, DESeq2 size-factor normalization and dispersion estimation were performed using default settings, and differential expression with the control group was assessed using a negative binomial generalized linear model with diet as the sole covariate. A Wald test for statistical significance was applied, and the Benjamini-Hochberg procedure was used to adjust for multiple comparisons. In parallel, single-cell differential expression was conducted using Wilcoxon rank-sum tests (Libra) comparing each high-fat diet group to control within cell type, using biological replicate as the unit of replication and restricting analyses to male mice and cell types with ≥50 cells per contrast.

